# Effect of vitamin C and E supplementation on human gastrointestinal tract tissues and cells: Raman spectroscopy and imaging studies

**DOI:** 10.1101/2021.11.04.467278

**Authors:** Krystian Miazek, Karolina Beton, Beata Brozek-Pluska

**Author notes:** Corresponding author: Beata Brozek-Pluska.

## Abstract

Cancer of gastrointestinal tract, such as colorectal cancer (CRC) and gastric cancer (GC), are common types of cancer globally and their origin can be linked to oxidative stress conditions. Commonly available antioxidants, such as vitamins C and E, are widely considered as potential anti-cancer agents. Raman spectra have great potential in the biochemical characterization of matter based on the fact that each molecule has its own unique vibrational properties. Raman spectroscopy allows to precisely characterized cell substructures (nucleus, mitochondria, cytoplasm, cell membrane) and components (proteins, lipids, nucleic acids).

The paper presents the application of the Raman spectroscopy technique for the analysis of tissue samples and cells of the human colon and stomach. The main goal of this study is to show the differences between healthy and cancerous tissues from the human digestive tract and human normal and cancer colon and gastric cell lines. The paper presents the spectroscopic characterization of normal colon cells - CCD-18 Co in physiological and oxidative conditions and effect of oxidative injury of normal colon cells upon supplementation with vitamin C at various concentrations based on Raman spectra. The obtained results were related to the Raman spectra recorded for human colon cancer cells - Caco-2. In addition, the effect of the antioxidant in the form of vitamin E on gastric cancer cells - HTB-135 is presented and compared with normal gastric cells - CRL-7869. All measured gastric samples were biochemically and structurally characterized by means of Raman spectroscopy and imaging. Statistically assisted analysis has shown that normal, ROS injured and cancerous human gastrointestinal cells can be distinguished based on their unique vibrational properties.

The conducted research based on Raman spectra proved that antioxidants in the form of vitamin C and E exhibit anti-cancer properties. In consequence, conducted studies proved that label-free Raman spectroscopy may play an important role in clinical diagnostics differentiation of human normal and cancerous gastrointestinal tissues and may be a source of intraoperative information supporting histopathological analysis.

## Introduction

Gastrointestinal tract cancer, a general term for a group of cancers localized within the digestive system and intestinal tract, includes colorectal cancer (CRC) and gastric cancer (GC) constituting a major public health problem worldwide.^1^

CRC, the third most common cancer in men and the second in woman worldwide, is characterized with high metastasis and poor prognosis.^2–4^ Development of CRC is correlated with aging, diet, lifestyle (smoking, alcohol consumption, physical inactivity) and genetic predispositions.^5,6^ Colon cancer originates from adenomatous polyps that progress into cancer or metastatic colon cancer.^7^ Strategies to prevent the occurrence and development of CRC comprise implementation of fecal occult blood tests, endoscopic surveillance of high-risk individuals for detection and removal of premalignant lesions.^6,8^

The majority of stomach cancer cases is adenocarcinoma, where cancer arises from epithelium in the mucosa, and can be divided into two types: intestinal and diffuse.^9^ The main reason of gastric cancer development is infection by bacterium *Helicobacter pylori*, although a style of life and genetic background are also important factors.^10,11^ The conventional diagnostic techniques for gastric cancer screening are incisional/excisional biopsy, upper endoscopy and contrast radiography.^12–14^

Another crucial strategy is the application of medications for successful anti-cancer therapy.^6,15^ Despite the ongoing development of synthetic anti-cancer drugs, there is still necessity to discover and investigate compounds available in nature, in terms of their chemotherapeutic and chemo-preventive effect against cancer occurrence and progression.^16^ Numerous molecules derived from plants and microbes, such as alkaloids, statins, polyphenols, isoprenoids, vitamins, can be considered for cancer therapy or prevention,^17,18^ vitamins: A, C, E are compounds that are worth being investigated.^19^ Vitamins are organic molecules involved in metabolic processes and cellular regulations, and are indispensable for growth and development.^20^ Vitamin C, is a representative of vitamins, that is widely discussed as a curing agent for prevention of many illnesses such as atherosclerosis, neurodegenerative disorders, diabetes and cancer.^21^ Vitamin C, also known as L-ascorbic acid (AA), performs a function as an antioxidant and free radical scavenger.^19^ Vitamin C is naturally present in fruits and vegetables,^21^ but can be also manufactured industrially.^22^ This vitamin is also a therapeutic agent for numerous diseases such as scurvy, colds and is considered as a therapeutic substance in the treatment of various type of cancer.^21^ Vitamin C was reported to exert anti-proliferative effect towards melanoma,^23^ adenocarcinoma gastric,^24^ breast cancer,^25^ pancreatic cancer,^26^ and colon cancer cells.^27^ Although ascorbic acid (AA) is a well-known antioxidant, it is said to react at high concentrations as a pro-oxidant and its anticancer effect is ascribed to its ability to induce autoxidation.^23,25^ Ascorbate, accumulated intracellularly, is autoxidized to dehydroascorbic acid (DHA), thereby providing electrons to oxygen and forming H_2_O_2_. Hence, elevated levels of reactive oxygen species (ROS) due to hydrogen peroxide formed, causes the toxicity to cancer cells. Induction of apoptosis,^27^ autophagy,^26^ impairment of mitochondrial structure,^28^ and Ca^2+^ release from mitochondria,^29^ was concluded to be a consequence of increase in ROS production, upon exposure to vitamin C in high doses.

Vitamin E consists of a family of 8 different molecules, 4 tocopherols (α-, β-, γ-, δ-) and 4 tocotrienols (α-, β-, γ-, δ-), each possessing a chromanol ring and a phytyl tail. Vitamin E are lipid soluble molecules that are localized in lipid regions of cell membranes. Vitamins E possess antioxidant properties as lipid peroxyl radical scavenger that donates the hydrogen ion from the phenol group on the chromanol ring, thereby preventing lipid peroxidation and preserving cellular membrane integrity.^30,31^ The natural sources of vitamin E are vegetables, fruits and nuts. α-Tocopherol is found in peanut, sunflower, cottonseed and safflower oil, and γ-tocopherol is predominantly found in corn, linseed, soybean and rapeseed oil.^32^ Tocopherols are considered to be potential molecules for the prevention of atherosclerosis, the progressive accumulation of lipid-loaded fibrous plaques within the artery wall, what leads to cardiovascular disorders.^30^ Vitamin E in different forms showed also anti-proliferative activity against various cancer cell lines.^33–37^ Interference in *de novo* sphingolipids synthesis,^33^ vacuole formation and chromatin condensation,^34,37^ reduction in DNA synthesis and fragmentation of DNA,^35^ apoptosis induction,^36,37^ was reported as a consequence of exposure to vitamin E.

Diagnostic techniques for detection of CRC and GC possess their own specific advantages and disadvantages in terms of their accessibility and accuracy of diagnostic outcomes.^8,14^ Therefore, it is necessary to investigate new techniques enabling rapid and accurate detection of cancer presence and development in the colon and stomach tissues. Particular attention has been put on testing spectroscopic techniques, such as Infrared Spectroscopy and Raman spectroscopy (RS), in terms of their application for medical diagnosis. RS is a non-destructive spectroscopic technique which aim is to analyse light scattered inelastically by vibrating molecules and to obtain information about the chemistry and structure of the investigated molecule/sample. A source of light, encountering a molecule/sample, are lasers emitting light of specific wavelengths such as e.g. 355 nm (UV), 532 nm (green), 633 nm (red) or 785 nm (near-infrared).^38,39^ RS was reported to find application for detection of skin diseases (atopic eczema, allergy, melasma),^40^ but also for different types of cancer (breast, lung, brain, pancreatic, prostate, ovarian, gastrointestinal),^39,41^ with successful differentiation between cancer and healthy tissues. Indeed, Raman spectroscopy and imaging can be also a useful tool for the structural characterization of the single cells, including brain,^42^ breast,^43^ and even colon,^44,45^ as well as the whole tissue areas.

Summarizing, occurrence of CRC and GC is a consequence of oxidative stress generation, which causes DNA damage leading to carcinogenesis.^3,46^ Vitamin C and E were reported to cause implications for metabolism and composition of the whole cells.^23–27 30–37^

In this study, the effect of vitamin C on the composition of ROS injured normal colon cells and the effect of vitamin E on the composition of gastric cancer cells are investigated. Raman spectroscopy and imaging are used for biochemical and structural characterization of normal colon cells, normal colon cells exposed to oxidative stress generated by Tert-butyl hydroperoxide (tBHP), normal colon cells exposed to oxidative stress and vitamin C treatment, as well as for gastric cells, and gastric cells exposed to vitamin E treatment. The influence of different concentrations of vitamin C under oxidative stress on chemical profile of colon cells, and different concentrations of vitamin E on chemical profile of gastric cells, is evaluated according to Raman spectra analysis of investigated cell lines. Additionally, Raman spectroscopy was used for chemical characterization of cancer and normal stomach tissues.

## Materials and methods

### Cell line and cultivation conditions

CCD-18 Co cell line (ATCC® CRL-1459 ™), used in this study was purchased from ATCC (The Global Bioresource Center). CCD-18 Co cells were cultured in ATCC-formulated Eagle’s Minimum Essential Medium with L-glutamine. For the complete growth medium, Fetal Bovine Serum was added to reach a final concentration of 10%. A fresh medium was used every 2–3 days. CaCo-2 cell line was also purchased from ATCC and cultured according to the ATCC protocols. The CaCo-2 cell line was obtained from a patient - a 72-year-old Caucasian male diagnosed with colon adenocarcinoma. The biological safety of the obtained material is clas-sified as level 1 (BSL - 1). To make the medium complete we based on Eagle’s Minimum Essential Medium with L-glutamine, with addition of a fetal bovine serum to a final concentration of 20%. The medium s renewed once or twice a week.

Hs 746T (HTB-135™) and Hs 738.St/Int (CRL-7869 ™) lines used in this study was also purchased from ATCC (The Global Bioresource Center) and were cultured in ATCC-formulated Dulbecco’s Modified Eagle’s Medium (DMEM) (ATCC: 30-2002™) with L-glutamine, glucose, sodium pyruvate and sodium bicarbonate. For the complete growth medium, fetal bovine serum was added to reach a final concentration of 10%. A fresh medium was used every 2–3 days. All cell lines were cultivated in flat-bottom flasks made of polystyrene, possessing a cell growth surface equal to 75 cm^2^. Flasks with culture were stored in an incubator under following conditions: 37 °C, 5% CO_2_, 95% air.

### Chemical compounds

L-Ascorbic acid, reagent grade, crystalline (catalogue Number: A7506-25G), Vitamin E, α-tocopherol (catalogue Number: 258024) were purchased from Merck Life Science Sp. z o.o. company, XTT proliferation Kit (catalogue Number: 20-300-1000) was purchased from Biological Industries. All chemical compounds were used without additional purification.

### Cell treatment with tBHP, vitamin C and vitamin E

CCD-18 Co and HTB-135 cells, seeded onto calcium fluoride (CaF_2_) windows (25×1 mm) at a density of 10^3^ cells/cm^3^, were incubated for 24 h. To induce oxidative stress conditions on CCD-18 Co cells standard growth medium was removed and 50 µM of tBHP solution (in grown medium) was added and subsequently 5, 25 or 50 µM solution of vitamin C confected in fresh culture medium was added for 24-h cultivation. For HTB-135 cells, 5, 25 or 50 µM of vitamin E solved in fresh culture medium was added for 24-h supplementation.

### Tissue sample preparation

Colon and stomach tissue samples were collected during routine surgery as an intraoperative biological material. The non-fixed, fresh samples were used to prepare 16 µm sections. Specimens of the tissue from the tumor mass and the tissue from the safety margins outside of the tumor mass were prepared for Raman analysis by placing specimens on CaF_2_ windows. Additionally, tissue sections were stained (hematoxylin and eosin) for standard histological analysis. The histopathological analysis was performed by pathologists from Medical University of Lodz, Department of Pathology, Chair of Oncology. All procedures performed in studies involving human participants were in accordance with the ethical standards of the institutional and/or national research committee and with the 1964 Helsinki Declaration. All tissue procedures were conducted under a protocol approved by the institutional Bioethical Committee at the Medical University of Lodz, Poland (RNN/323/17/KE/17/10/2017, 17, October 2017). Written informed consent was obtained from patients.

### Raman spectroscopy and imaging of cells and tissues

After 24h of supplementation, cells were washed with Phosphate Buffered Saline (PBS, Gibco, catalogue Number: 10010023, pH 7.4 at 25°C, 0.01 M) to remove any residual medium components and an excess of additives that did not penetrate inside the cells during supplementation. Furthermore, PBS was removed and cells were fixed in 4% buffered formaldehyde solution for 10 min, and rinsed again with PBS. Prepared cells samples and tissue sections on CaF_2_ windows were subjected to Raman spectroscopy measurements.

Raman spectra and images of cells were registered using the confocal microscope Alpha 300 RSA+ (WITec, Ulm, Germany) equipped with an Olympus microscope integrated with a fiber (50 µm core diameter), with a UHTS spectrometer (Ultra High Through Spectrometer) and a CCD Andor Newton DU970NUVB-353 camera operating in default mode at −60 °C in full vertical binning mode. Nikon objective lens, with 40x magnification and a numerical aperture (NA = 1.0), was used for cell measurements carried out by immersion in PBS, for tissues samples measurements 40x dry objective (Nikon, objective type CFI Plan Fluor C ELWD DIC-M, numerical aperture (NA) of 0.60 and a 3.6–2.8 mm working distance) was used. During the experiments, a 532 nm excitation line, with the excitation laser power of 10 mW, with an integration time of 0,3s for Raman measurements for the high frequency region and 0,5 s for the low frequency region was used. All data for colon and stomach cells and colon tissues was collected and processed with the use of WITec Project Plus software. All spectroscopic and imaging data were analyzed by Cluster Analysis (CA), executed using WITec Project Plus software with Centroid model and k-means algorithm. Data normalization was performed using the normalization model by means of the Origin software.

Raman spectra and images of stomach tissues were investigated under Renishaw inVia™ confocal Raman microscope (New Mills Wotton-under-Edge, United Kingdom) using a ion laser (Ar) with excitation laser line 532 nm. All experiments were carried out at room temperature. The measurement parameters included grating at 600 l/mm, laser excitation power at 50%, which is equal to 12.2 mW, and exposure time set at 0.5/s. Optical microscope (Leica) equipped with an objective (Nikon, Plan Fluor) with 40x magnification, was used to visualize the morphology of the tested sample. The same parameters were applied whilst measuring each sample. Mapping was carried out to record chemical images (resolution 1 μm) of tissue areas. Average spectrum was obtained from each recorded image. Computations were performed with the use of WiRE™ software, where obtained spectra were proceeded with truncating, cosmic ray removal, noise filtering and baseline subtracting. Moreover, single spectra, proceeded with Cluster Analysis method, were grouped into clusters represented by centroids.

### Cluster analysis

Spectroscopic data were analyzed using Cluster Analysis method. This method constitute an exploratory form of comparative data analysis in which the observations are divided into different groups that share a set of common features - in this report it is related to vibrational characteristics. Cluster Analysis was made using WITec Project Plus software (colon tissues, colon cells, gastric cells) or Renishaw WiRE™ (stomach tissues) software with Centroid model and k-means algorithm, where each cluster is represented by one mean vector.

Cluster Analysis constructs groups (or classes or clusters) based on the principle that, within a group, the observations must be as similar as possible, while observations belonging to different groups must be different. The partition of n observations (x) into k (k≤n) clusters S should be done to minimize the variance (Var) according to the formula:

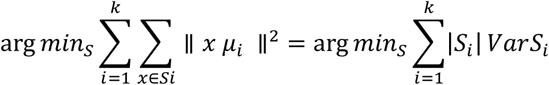

where *μ*_*i*_ is the mean of points.

All Raman maps presented in this research work were constructed based on principles of Cluster Analysis described above. Number of clusters (the minimum number of clusters characterized by different average Raman spectra, which describe the variety of the inhomogeneous biological sample) was 7 and 2 or 3 for cells and tissues: normal and cancerous, respectively.

### Data processing

Data acquisition and processing were performed using WITec Project Plus or Renishaw WiRETM software. Cosmic rays were removed from each Raman spectrum and the Savitzky-Golay method was implemented for the smoothing procedure. The background subtraction and the normalization (model: divided by norm (divide the spectrum by the dataset norm)) were performed by using Origin software. The normalization model: divided by norm was performed according to the formula:

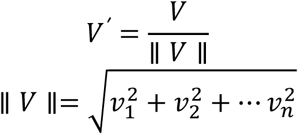

where: ν_*n*_ is the n^th^ V values.

The normalization was performed for the spectral region: 500-1800 cm^−1^, which forms a peculiar fingerprint of the sample.

### Statistical analysis

All results regarding the analysis of the intensity of the Raman spectra as a function of the type and time supplementation are presented as the mean ± SD, p < 0.05; where: SD - standard deviation, p – probability value. ANOVA analysis (Analysis of variance) was conducted using Origin software (significance level – 0.05, range test-Tukey).

### XTT cell viability assay

Cells were seeded at 5×10^3^ cells/well in a 96-well plate and incubated overnight. For experiments with CCD-18 Co cells, they were treated with 50 µM solution of tBHP - Reactive Oxygen Species generating agent and then various concentrations of vitamin C were added. After 24-h supplementation in standard cultivation conditions, cells were subjected to cell viability test with tetrazolium salts (XTT).

Colorimetric assays analyze the number of viable cells by the cleavage of tetrazolium salts added to the culture medium. This technique requires neither washing nor harvesting of cells, and the complete assay, from microculture to data analysis by an BioTek Synergy HT reader, is performed in the same microplate.

Tetrazolium salt is cleaved to formazan by a complex cellular mechanism. This bioreduction occurs in viable cells only, and is related to NAD(P)H production through glycolysis. Therefore, the amount of formazan dye formed directly correlates to the number of metabolically active cells in the culture. The intensity of the coloured product is directly proportional to the number of viable cells present in the tested culture. The percentage inhibition in each assay was calculated and plotted. tBHP treatment can lead to cell DNA damage followed by cell cycle arrest.

The assay determines the metabolic activity of living cells converting tetrazolium salts to formazan, which is measured by colorimetric tests at 450 nm wavelength, from which the specific signal of the sample is obtained, and 650 nm wavelength, which is the referential sample during the test on a multi-detecting BioTek Synergy HT model reader.

The effect of tBHP and vitamin C on viability of CCD-18 Co cells was investigated. Results show (Scheme 1) that tBHP can decrease viability of CCD-18 Co culture, but addition of different vitamin C concentrations can greatly improve viability of this cells.

Scheme 1 shows the results of XTT test obtained for CCD-18Co human normal colon cells supplemented with ROS generating agent at concentration 50 µM and vitamin C in various concentrations after 24h supplementation.

**Scheme 1.**
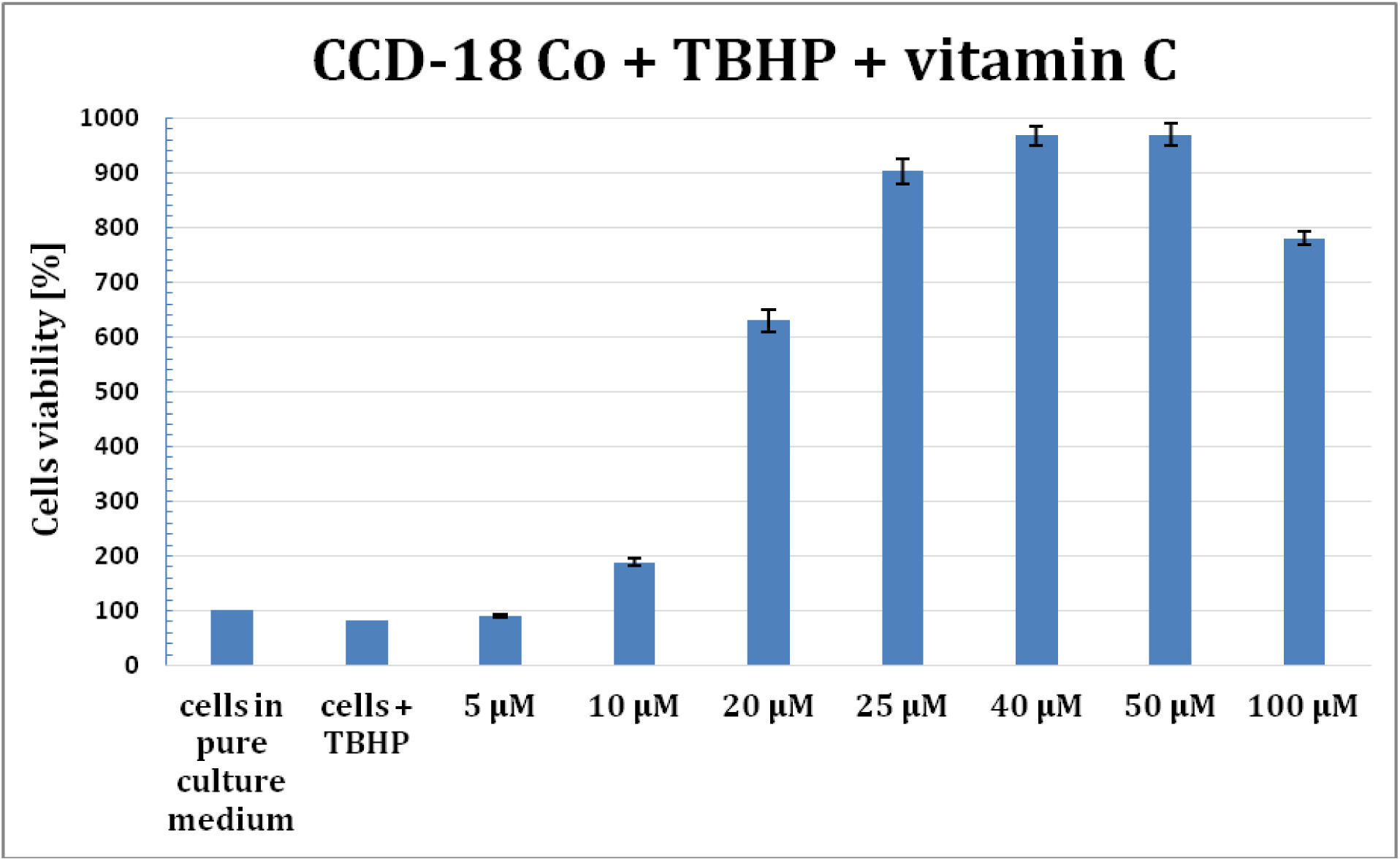
Results of XTT comparison of the percent viability for CCD-18 Co human normal colon cells treated with tBHP and supplemented with different concentrations of vitamin C after 24h supplementation, mean±SD, SD-standard deviation.

The results of measurements carried out for CCD-18 Co cells with the addition of the stress compound tBHP showed that under oxidative stress conditions, the growth and development of cells with normal structure is disturbed and is initially inhibited in such a way that the cells stop at the stage they were at when they appeared stressful conditions. With prolonged exposure to unfavourable, oxidative conditions, their survival gradually begins to decline, and the weaker or less developed ones probably die already at this stage. This phenomenon is illustrated by the bar of the 24-hour incubation with tBHP, for which the viability of the cells is lower than for the control sample. The attached data show that a higher concentration of the antioxidant vitamin C results in a faster increase in cell survival and proliferation when subjected to oxidative conditions. Vitamin C reduces the harmful stress effect caused by ROS. Based on XTT tests results concentrations of vitamin C equal to 5, 25 and 50 µM were chosen for spectroscopic experiments. The same concentrations were chosen for vitamin E supplementations for experiments for gastric human cancer cells.

## Results

In performed spectroscopic analysis, Raman spectroscopy and imaging were used for biochemical characterization of CCD-18 Co cells, CCD-18 Co cells treated with tBHP and different vitamin C concentrations, HTB-135 cells, HTB-135 cells treated with different vitamin E concentrations (but the same concentrations as for vitamin C), and Caco-2 and CRL-7869 cells as reference cell samples of cancerous colon cell line and normal stomach cell line, respectively.

In general, Raman vibrational spectra consists of two interesting regions: the Raman fingerprint region: 500–1800 cm^−1^ and the high frequency region: 2700–3100 cm^−1^ (the region 1800-2700 cm^−1^ is excluded from consideration through the lack of Raman bands). In this manuscript we focused on most informative fingerprint region 500–1800 cm^−1^.

To properly rise to biochemical changes in both normal and cancerous human colon and stomach cell lines and tissues by Raman spectroscopy and imaging, we will closely inquire how the Raman based method responds to generated ROS and changes related to vitamins supplementation. The experiments will extend our knowledge of the protective function of antioxidants and veritable influence of the ROS generation on cancer development by RS.

All Raman spectra contain bands assigned to specific chemical structures, based on vibrational features of molecules within an analysed cell. Table 1 featured the main chemical constituents which can be identified based on their vibrational properties in analysed gastrointestinal human tissues, and human colon and gastric cells in normal and oxidative stress conditions generated by tBHP adding (discussed later in the manuscript), and upon vitamin C and vitamin E supplementation.

**Table 1.** The tentative assignments of Raman peaks for chemical characterization of CCD-18 Co (normal colon), Caco-2 (cancer colon), CRL-7869 (normal gastric), and HTB-135 (cancer gastric) cells used in this study.^47^

| Wavenumber [$\text{cm}^{-1}$] | Tentative Assignments |
| --- | --- |
| 716-723 | DNA (nucleotides) |
| 748-757 | Tryptophan (protein assignment) |
| 781-787 | DNA (nucleotides) |
| 852-858 | Proline, hydroxyproline, tyrosine |
| 992-1010 | Phenylalanine |
| 1070-1093 | Phosphodiester groups in nucleic acids and phospholipids |
| 1127-1133 | Fatty acids, lipids |
| 1256-1268 | Amide III band in proteins |
| 1299-1305 | Fatty acids, lipids, phospholipids |
| 1337-1348 | Proteins, nucleic acids |
| 1444-1447 | Lipids and proteins |
| 1582-1854 | Phosphorylated proteins |
| 1658-1664 | Proteins (amide I), lipids, nucleic acids |

Spectroscopic analysis by Raman spectroscopy and imaging was started with gastrointestinal samples without any supplementation.

Fig. 1 characterize single, human normal colon CCD-18 Co cell by depicting the microscopic image, Raman image of single cell constructed based on Cluster Analysis (CA) method, Raman images of all clusters identified by CA assigned to: nucleus, mitochondria, lipid-rich regions, cytoplasm, membrane, and cell environment, and representative average spectra from fingerprint region (500-1800 cm^−1^).

**Fig. 1.**
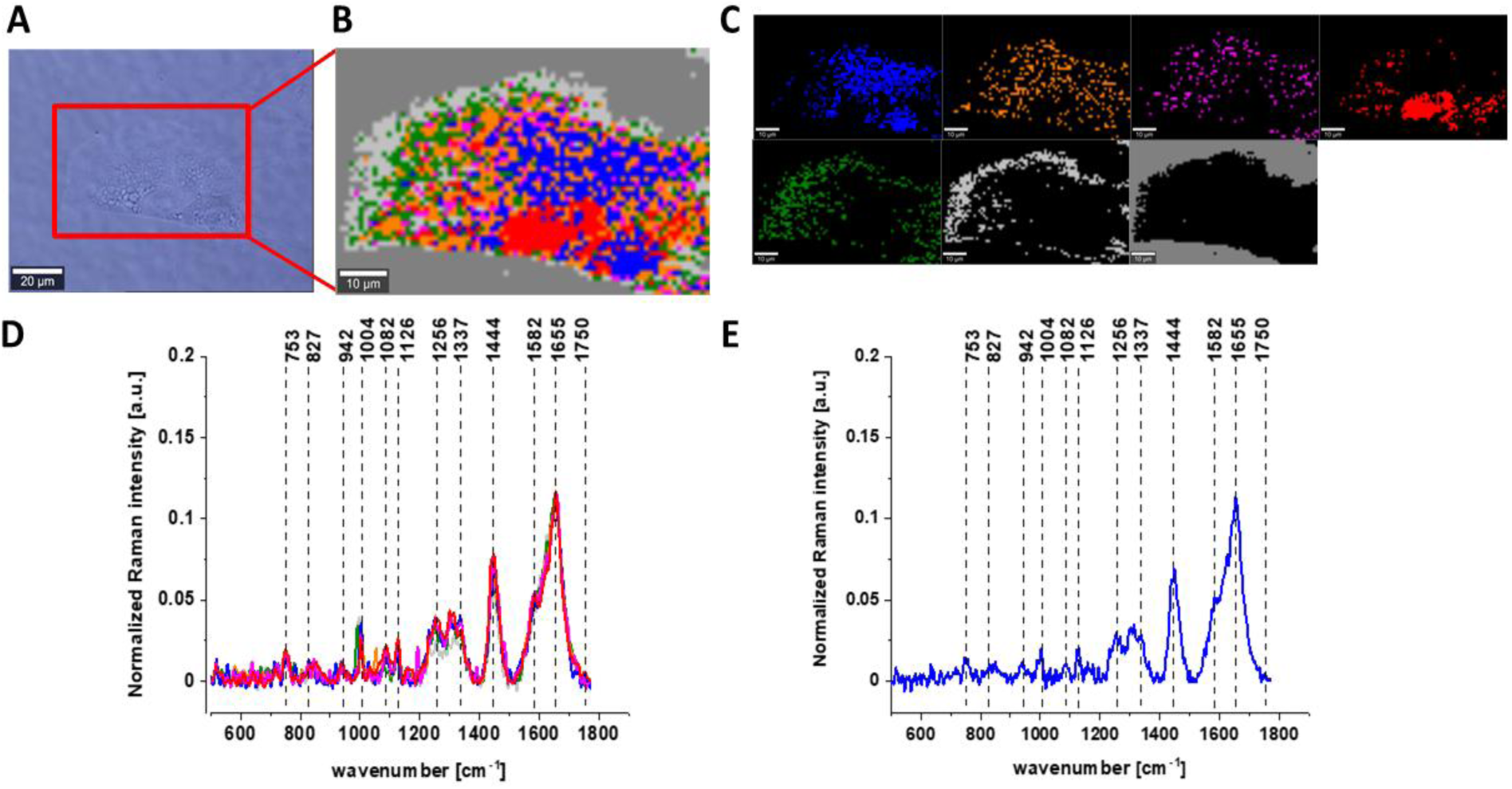
The microscopy image of normal colon CCD-18 Co cell (A), Raman image constructed based on Cluster Analysis (CA) method (B), Raman images of all clusters identified by CA assigned to: nucleus (red), mitochondria (magenta), lipid-rich regions (blue, orange), cytoplasm (green), membrane (light grey), and cell environment (dark grey) (C), average Raman spectra typical for all clusters identified by CA in a fingerprint region (D), average Raman spectrum for the whole cell in a fingerprint region (E), cells measured in PBS, excitation wavelength 532 nm. Adapted with permissions.^48^

**Fig. 2.**
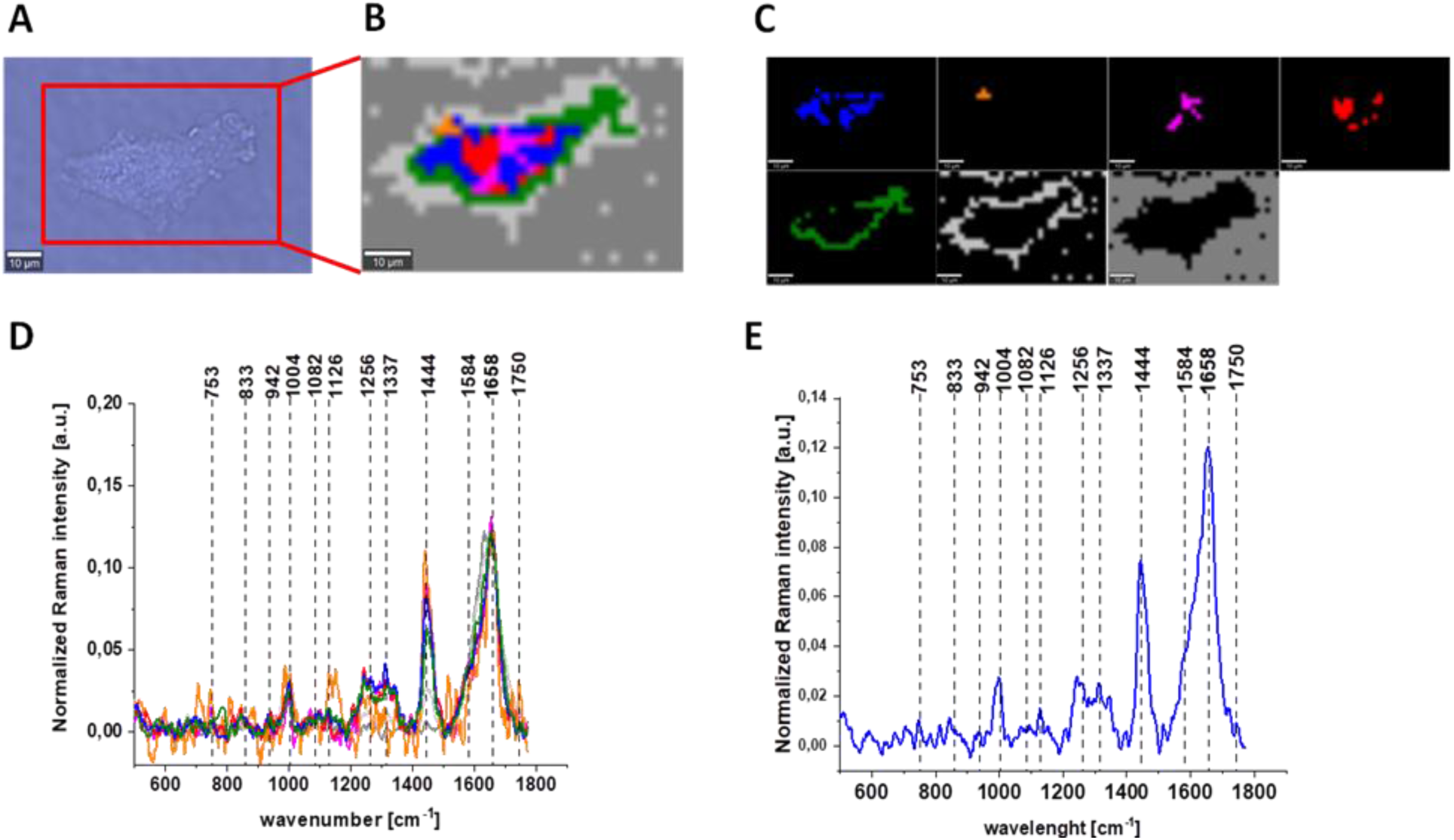
The microscopy image (A), Raman image constructed based on Cluster Analysis (CA) method (B) of normal colon cell CCD-18 Co treated with tBHP at concentration 50 µM, Raman images of all clusters identified by CA assigned to: nucleus (red), mitochondria (magenta), lipid-rich regions (blue, orange), cytoplasm (green), membrane (light grey), and cell environment (dark grey) (C), average Raman spectra typical for all clusters identified by CA in a fingerprint region (D), average Raman spectrum for the whole cell in a fingerprint region (E), cells measured in PBS, excitation wavelength 532 nm.

Results for CCD-18 Co cells treated with tBHP, and 5, 25 and 50 µM of vitamin C upon ROS generation, and Caco-2 cells are also presented below.

Raman bands in spectra presented in Figs. 1-6 on panels D are ascribed to nucleic acids, amino acids, proteins and lipids, thereby providing chemical characterization of CCD-18 Co and Caco-2 cells (see Table 1).

One of the major goal of the undertaken research was the biochemical analysis of human normal colon cell line in physiological or oxidative stress conditions, demonstrating the antioxidant properties of vitamin C and comparative analysis of normal human colon cells in different conditions with human cancerous colon cells Caco-2 based on the vibrational features by using label-free Raman spectroscopy and imaging. Fig. 3 shows Raman spectra and imaging for cancerous human colon cells Caco-2.

**Fig. 3.**
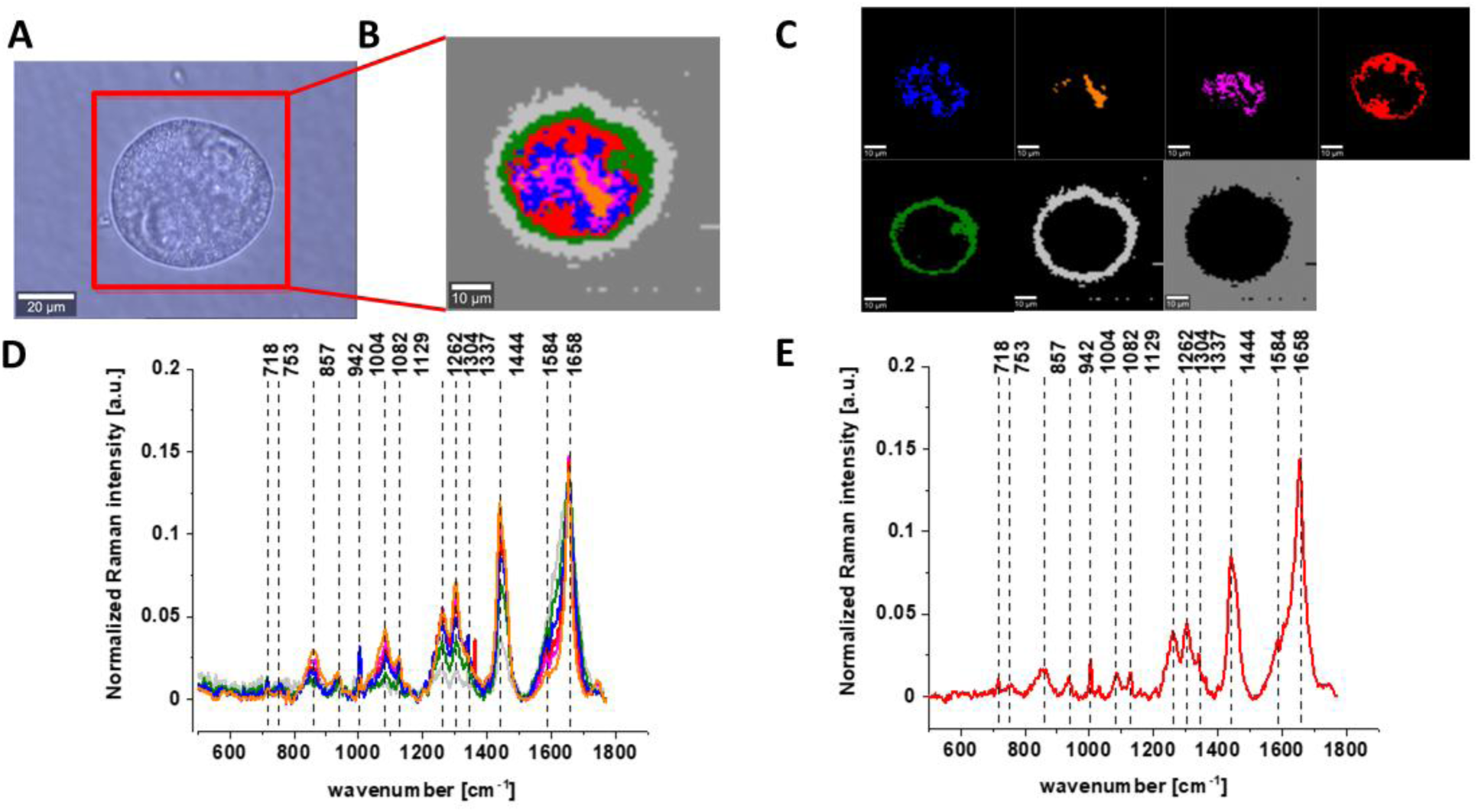
The microscopy image of colon cancer Caco-2 cells (A), Raman image constructed based on Cluster Analysis (CA) method (B), Raman images of all clusters identified by CA assigned to: nucleus (red), mitochondria (magenta), lipid-rich regions (blue, orange), cytoplasm (green), membrane (light grey), and cell environment (dark grey) (C), average Raman spectra typical for all clusters identified by CA in a fingerprint region (D), average Raman spectrum for the whole cell in a fingerprint region (E), cells measured in PBS, excitation wavelength 532 nm. Adapted with permissions.^45^

One can see from Figs. 1-3 that using Raman spectroscopy it is possible to obtain well resolved vibrational spectra to characterize the biochemistry of single cells.

Figs. 4-6 present microscopy and Raman data obtained for CCD-18 Co human normal colon cells treated by using tBHP and upon vitamin C supplementation with different concentrations.

**Fig. 4.**
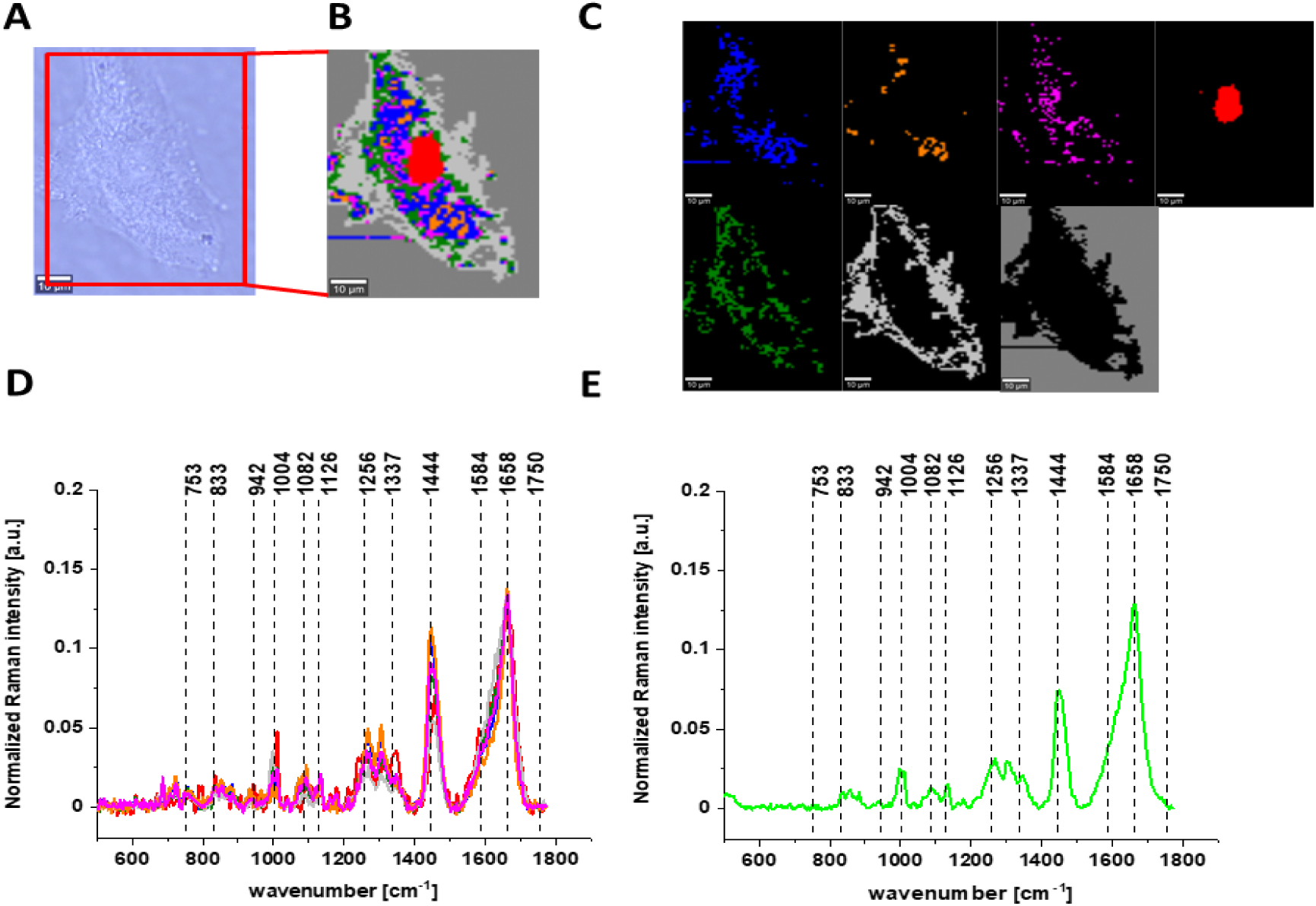
The microscopy image of normal colon CCD-18 Co cell subjected to vitamin C (5 µM) and tBHP (50 µM) treatment (A), Raman image constructed based on Cluster Analysis (CA) method (B), Raman images of all clusters identified by CA assigned to: nucleus (red), mitochondria (magenta), lipid-rich regions (blue, orange), cytoplasm (green), membrane (light grey), and cell environment (dark grey) (C), average Raman spectra typical for all clusters identified by CA in a fingerprint region (D), average Raman spectrum for the whole cell in a fingerprint region (E), cells measured in PBS, excitation wavelength 532 nm.

**Fig. 5.**
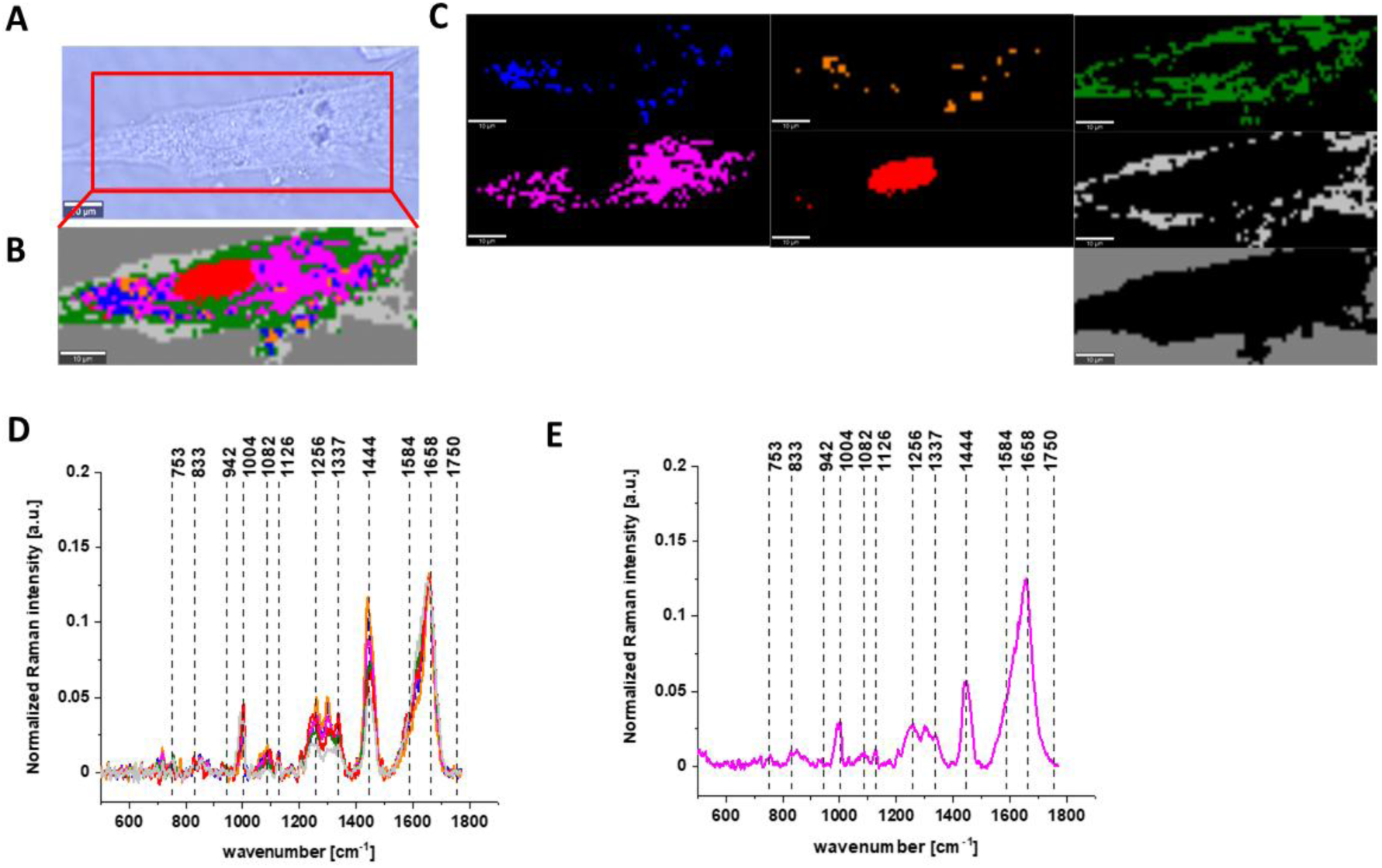
The microscopy image of normal colon CCD-18 Co cell subjected to vitamin C (25 µM) and tBHP (50 µM) treatment (A), Raman image constructed based on Cluster Analysis (CA) method (B), Raman images of all clusters identified by CA assigned to: nucleus (red), mitochondria (magenta), lipid-rich regions (blue, orange), cytoplasm (green), membrane (light grey), and cell environment (dark grey) (C), average Raman spectra typical for all clusters identified by CA in a fingerprint region (D), average Raman spectrum for the whole cell in a fingerprint region (E), cells measured in PBS, excitation wavelength 532 nm.

**Fig. 6.**
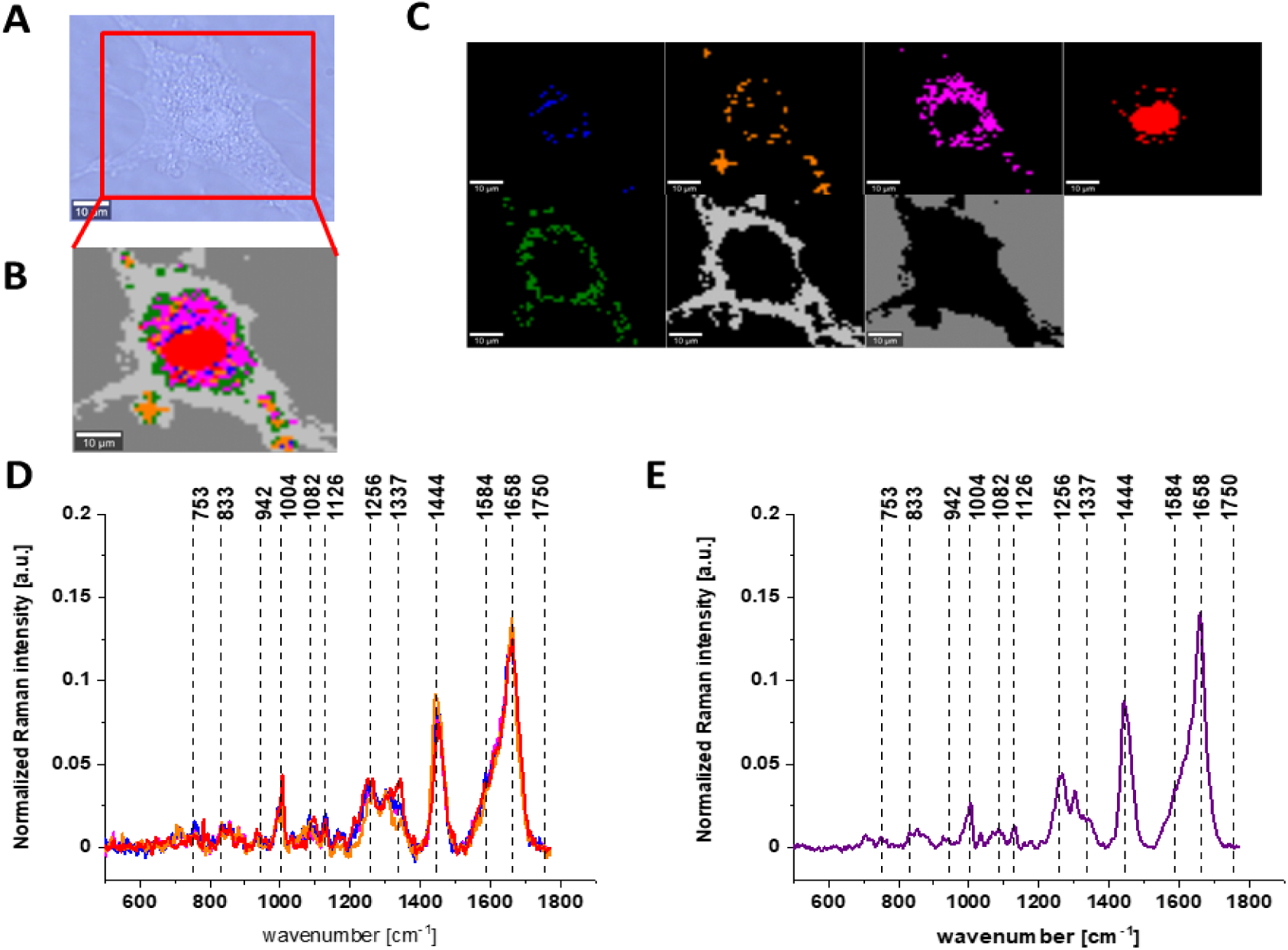
The microscopy image of normal colon CCD-18 Co cells subjected to vitamin C (50 µM) and tBHP (50 µM) treatment (A), Raman image constructed based on Cluster Analysis (CA) method (B), Raman images of all clusters identified by CA assigned to: nucleus (red), mitochondria (magenta), lipid-rich regions (blue, orange), cytoplasm (green), membrane (light grey), and cell environment (dark grey) (C), average Raman spectra typical for all clusters identified by CA in a fingerprint region (D), average Raman spectrum for the whole cell in a fingerprint region (E), cells measured in PBS, excitation wavelength 532 nm.

The same spectroscopic analysis by using Raman spectroscopy and imaging was performed for human normal and cancer gastric cells CRL-7869 and HTB-135 respectively.

Figs. 7-11 characterize single HTB-135 and CRL-7869 cells by depicting the microscopy images, Raman images of single cells constructed based on Cluster Analysis (CA) method, Raman images of all clusters identified by CA assigned to: nucleus, mitochondria, lipid-rich regions, cytoplasm, membrane, and cell surroundings, and representative cluster spectra from fingerprint region (500-1800 cm^−1^).

**Fig. 7.**
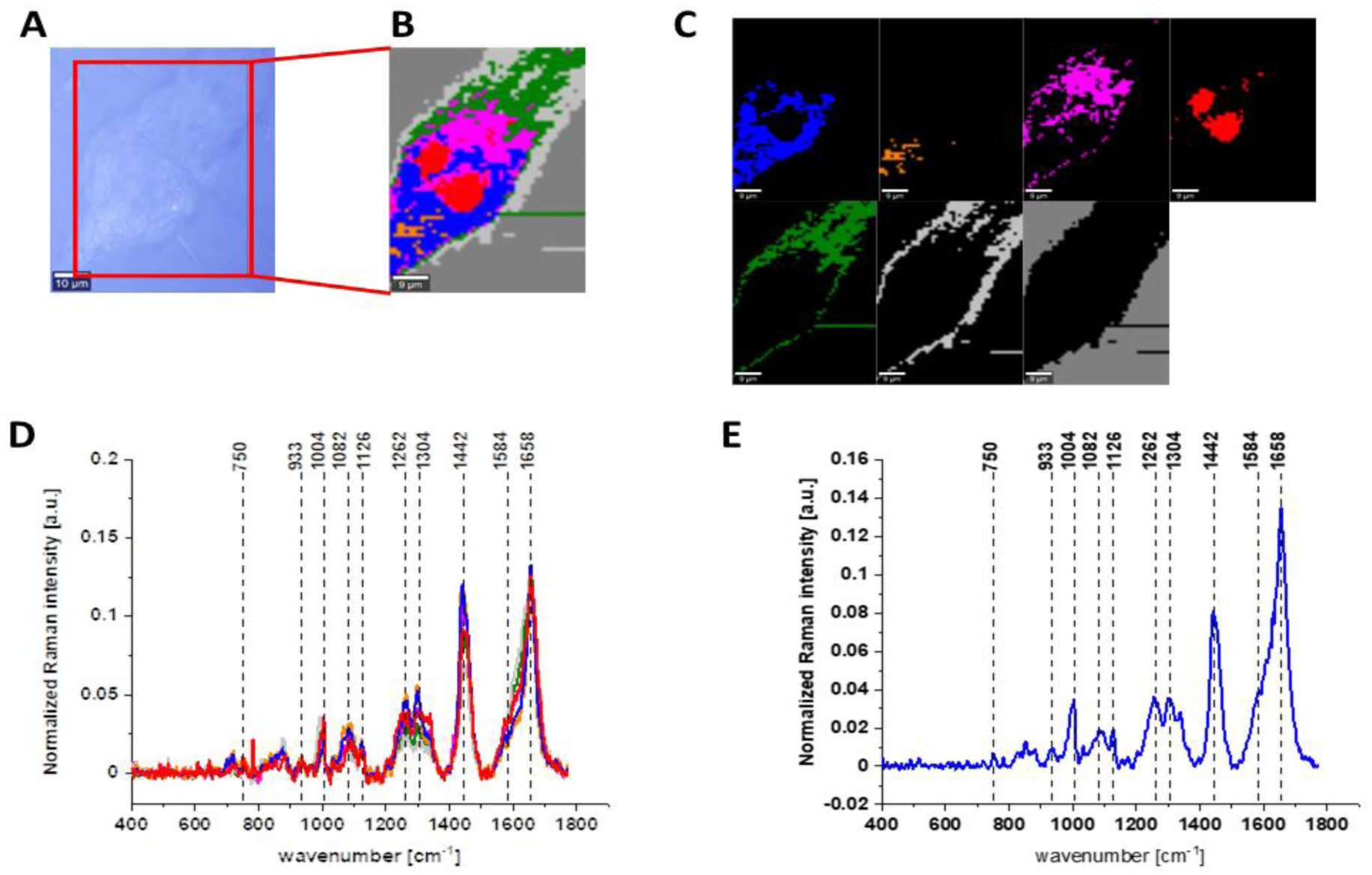
The microscopy image of human normal gastric CRL-7869 cell (A), Raman image constructed based on Cluster Analysis (CA) method (B), Raman images of all clusters identified by CA assigned to: nucleus (red), mitochondria (magenta), lipid-rich regions (blue, orange), cytoplasm (green), membrane (light grey), and cell environment (dark grey) (C), average Raman spectra typical for all clusters identified by CA in a fingerprint region (D), average Raman spectrum for the whole cell in a fingerprint region (E), cells measured in PBS, excitation wavelength 532 nm.

Results for HTB-135 cancer gastric cells treated with 5, 25 and 50 µM of vitamin E, and normal gastric CRL-7869 are also presented. Signals in spectra presented in Figs. 7-11 are ascribed to nucleic acids, amino acids, proteins and/or lipids, thereby providing biochemical characterization of HTB-135 and CRL-7869 cells (see Table 1).

Fig. 7 shows Raman spectra and Raman imaging for untreated normal gastric cells CRL-6879 measured in PBS.

Fig. 8 shows the results of the same type of spectroscopic analysis performed for gastric cancer cells HTB-135.

**Fig. 8.**
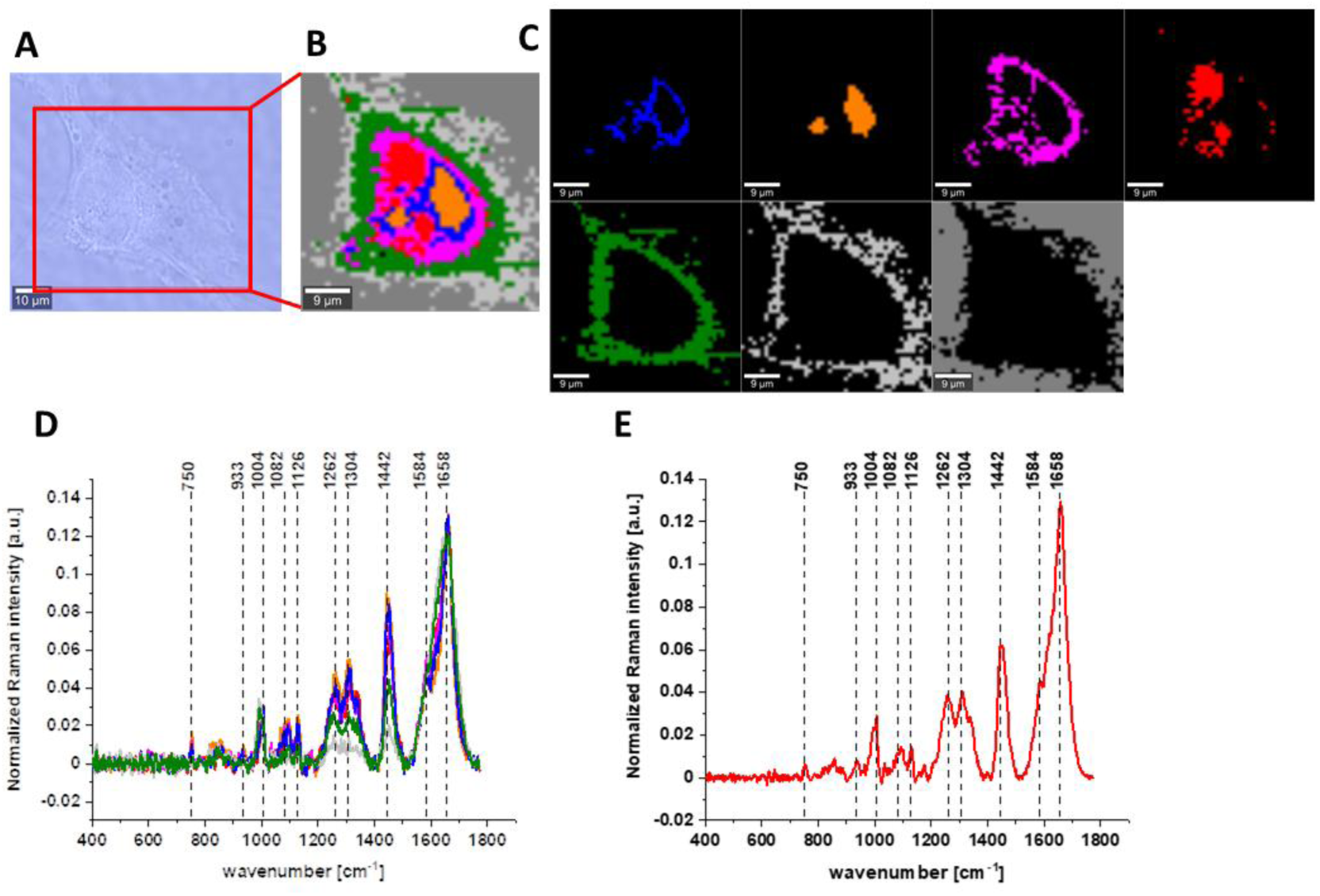
The microscopy image of human gastric cancer HTB-135 cell (A), Raman image constructed based on Cluster Analysis (CA) method (B), Raman images of all clusters identified by CA assigned to: nucleus (red), mitochondria (magenta), lipid-rich regions (blue, orange), cytoplasm (green), membrane (light grey), and cell environment (dark grey) (C), average Raman spectra typical for all clusters identified by CA in a fingerprint region (D), average Raman spectrum for the whole cell in a fingerprint region (E), cells measured in PBS, excitation wavelength 532 nm.

Figs. 9-11 present microscopy and Raman data obtained for HTB-135 human cancer gastric cells upon vitamin E supplementation with different concentrations: 5, 25 and 50 µM.

**Fig. 9.**
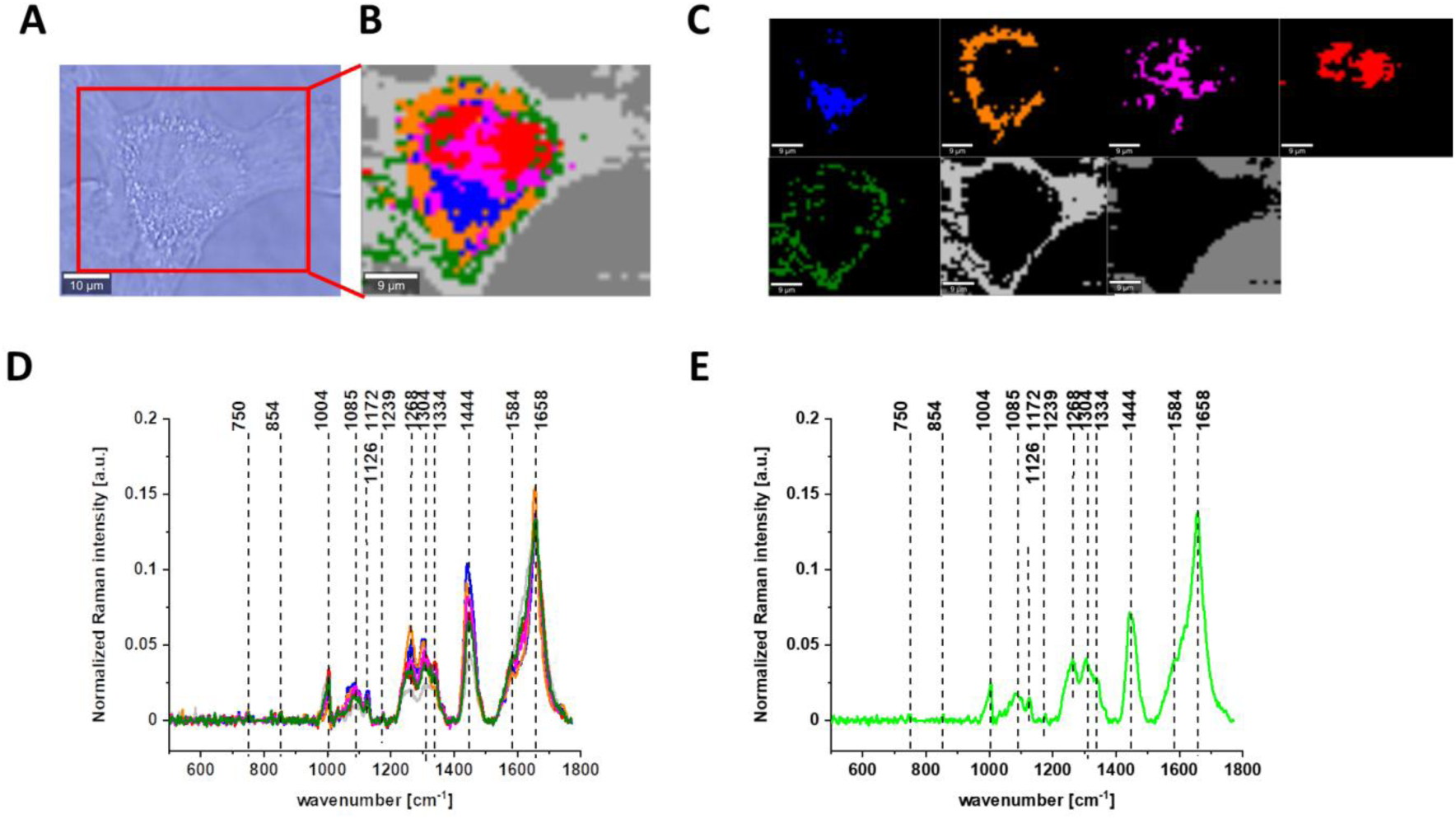
The microscopy image of human cancer gastric HTB-135 cell exposed to 5 µM vitamin E (A), Raman image constructed based on Cluster Analysis (CA) method (B), Raman images of all clusters identified by CA assigned to: nucleus (red), mitochondria (magenta), lipid-rich regions (blue, orange), cytoplasm (green), membrane (light grey) and cell environment (dark grey) (C), average Raman spectra typical for all clusters identified by CA in a fingerprint region (D), average Raman spectrum for the whole cell in a fingerprint region (E), cells measured in PBS, excitation wavelength 532 nm.

**Fig. 10.**
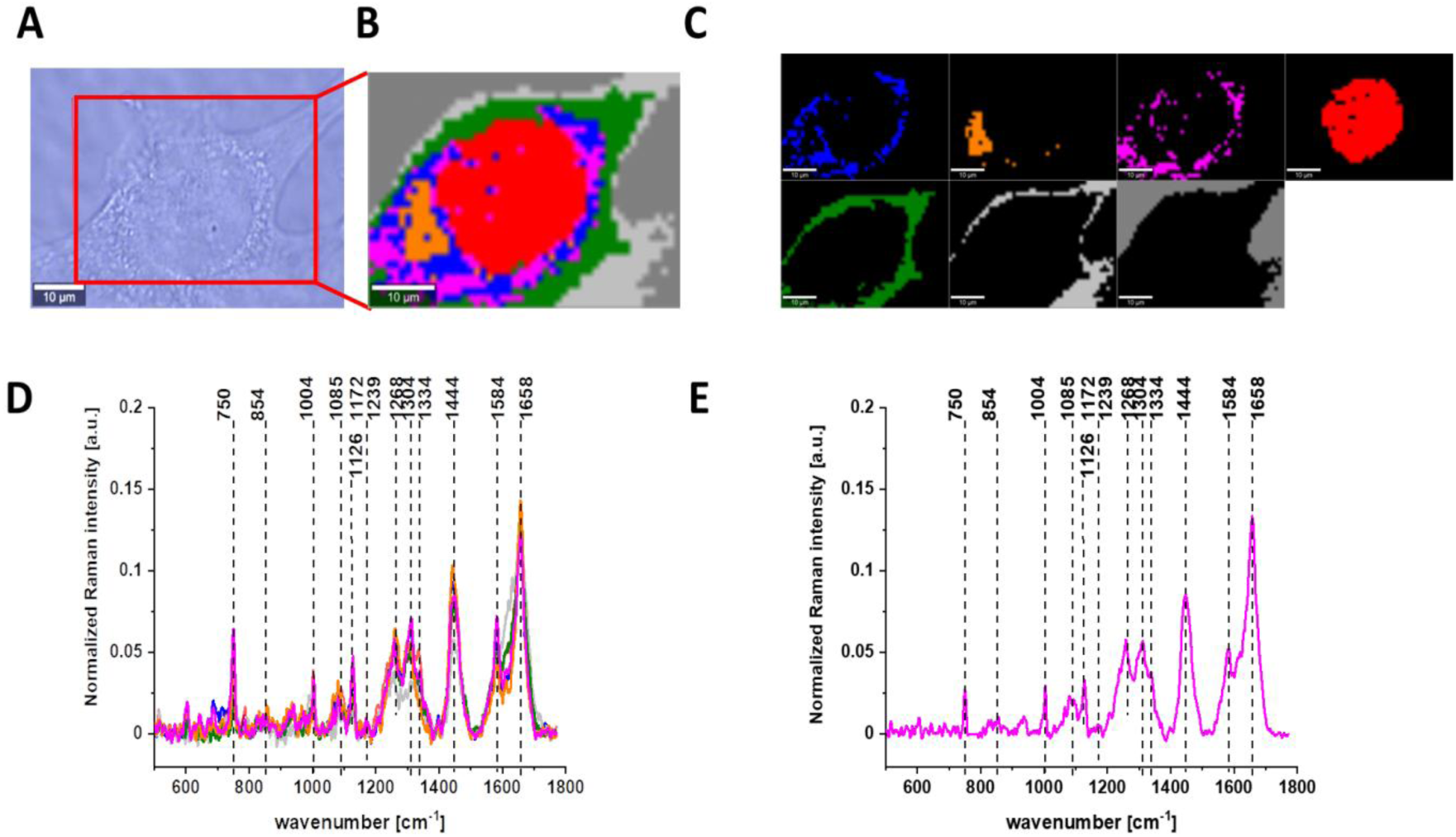
The microscopy image of human cancer gastric HTB-135 cell exposed to 25 µM vitamin E (A), Raman image constructed based on Cluster Analysis (CA) method (B), Raman images of all clusters identified by CA assigned to: nucleus (red), mitochondria (magenta), lipid-rich regions (blue, orange), cytoplasm (green), membrane (light grey), and cell environment (dark grey) (C), average Raman spectra typical for all clusters identified by CA in a fingerprint region (D), average Raman spectrum for the whole cell in a fingerprint region (E), cells measured in PBS, excitation wavelength 532 nm.

The results obtained by microscopy and spectroscopic techniques for human normal and cancer cells of colon and stomach were completed with data recorded for healthy and cancer human colon and stomach tissues.

Fig. 12 shows the data obtained by Raman spectroscopy and imaging of human normal and cancer colon tissues. As for single cells Raman spectra are characterize by peaks corresponding for nucleic acids, proteins and lipids including unsaturated fraction.

**Fig. 11.**
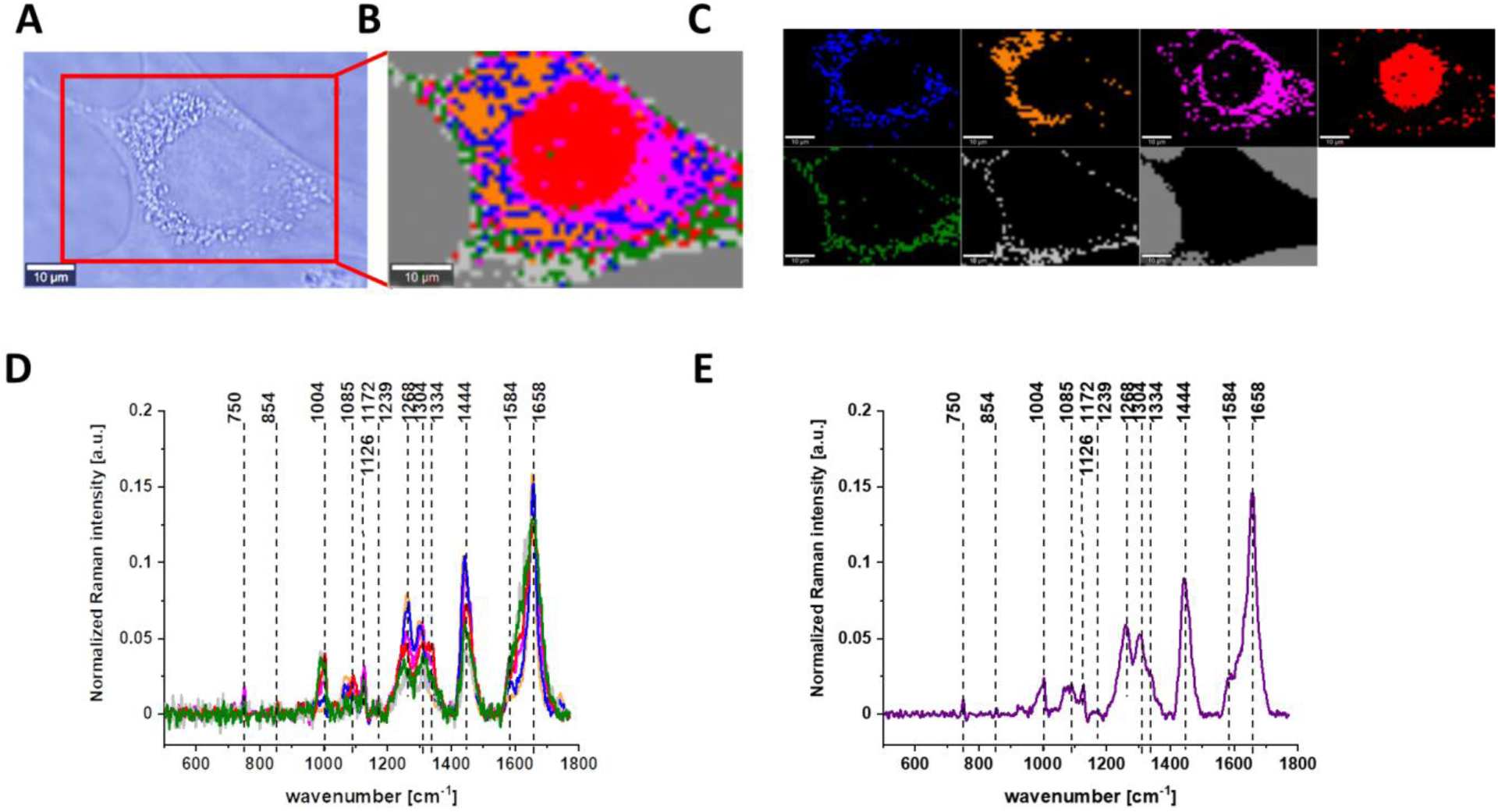
The microscopy image of human cancer gastric HTB-135 cell exposed to 50 µM vitamin E (A), Raman image constructed based on Cluster Analysis (CA) method (B), Raman images of all clusters identified by CA assigned to: nucleus (red), mitochondria (magenta), lipid-rich regions (blue, orange), cytoplasm (green), membrane (light grey), and cell environment (dark grey) (C), average Raman spectra typical for all clusters identified by CA in a fingerprint region (D), average Raman spectrum for the whole cell in a fingerprint region (E), cells measured in PBS, excitation wavelength 532 nm.

**Fig. 12.**
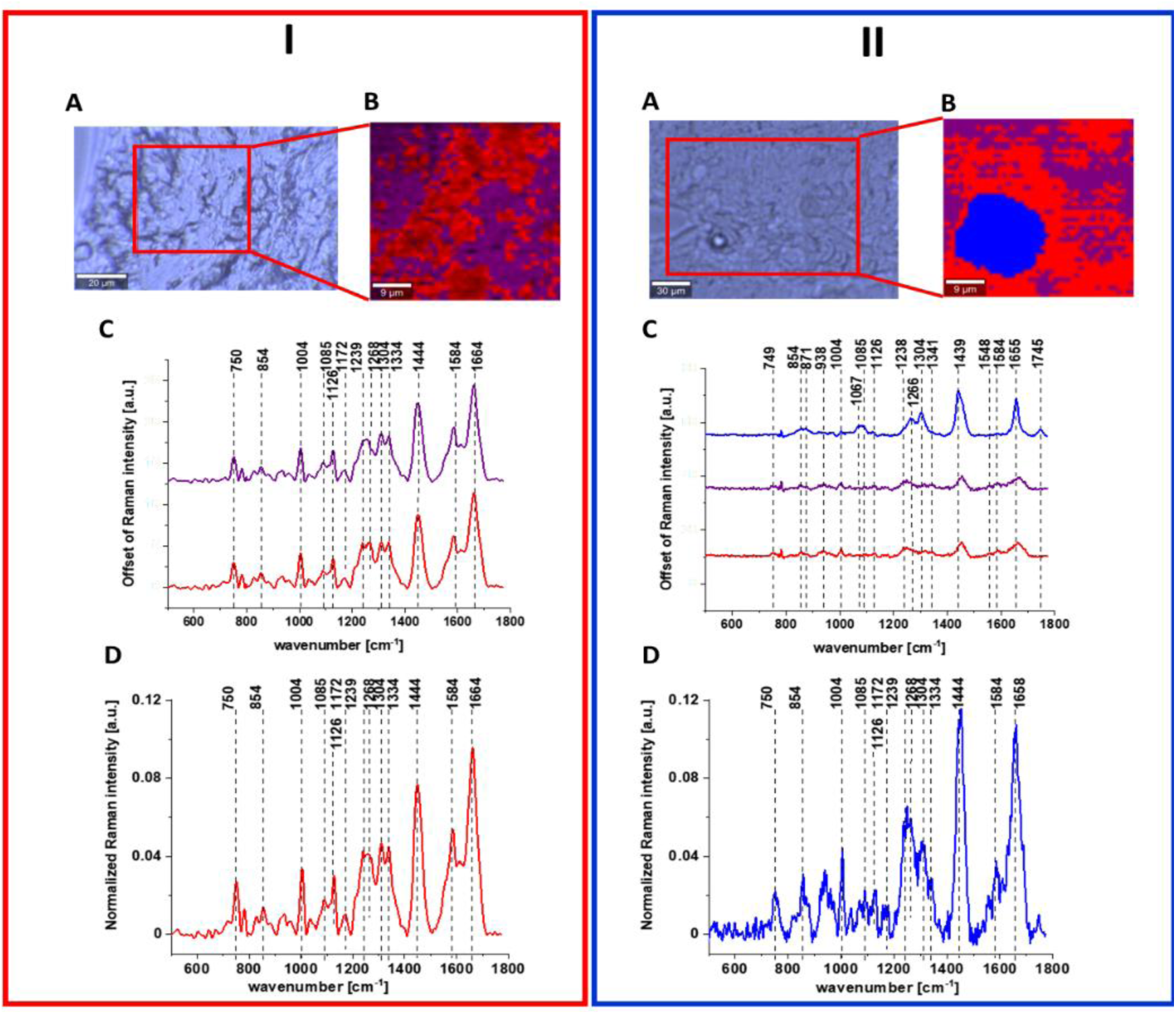
Raman image and spectral analysis of human colon cancerous tissue (I) and human colon normal tissue (II). The microscopy image (A), Raman image constructed by CA method (B), average Raman spectra typical of all clusters, colors of the spectra correspond to colors of clusters seen in B (C), and average (arithmetic mean) Raman spectrum for the entire area of analyzed tissue (D), excitation wavelength 532 nm. Adapted with permissions.^45^

Detailed analysis of data presented in Figs. 1-12 shows that Raman spectra typical for human colon cells and tissues are characterize by bands at c.a. 750 cm^−1^ (nucleic acids, DNA, tryptophan, nucleoproteins), c.a. 854 cm^−1^ (phosphate groups), 1004 cm^−1^ (phenylalanine), 1085 cm^−1^ (phosphodiester groups in nucleic acids), 1126 cm^−1^ (saturated fatty acids), 1172 cm^−1^ (cytosine, guanine), 1239 cm^−1^ (nucleic acids, Amide III), 1268 cm^−1^ (Amide III (C–N stretch + N–H bend)), 1304 cm^−1^ (lipids), 1334 cm^−1^ (CH_3_CH_2_ wagging vibrations of collagen), 1444 cm^−1^ (lipids (predominantly) and proteins), 1584 cm^−1^ (phosphorylated proteins) and 1664 cm^−1^ (Amide I (C=O stretch)) confirming that chemical information about the composition of cancerous and normal human colon cells and tissues can be obtained based on vibrational features of samples.

Moreover, Raman imaging allow not only to characterize colon samples quantitatively, but also provide information about spatial distribution of chemical compounds in analysed specimens and single cells.

Raman imaging study and Raman cluster analysis of colon tissues have shown also higher structural heterogeneity of normal tissue in a form of 3 clusters, and higher homogeneity of cancer tissue in a form of 2 clusters, indicating that colon cancerous transformations cause more uniform alterations during pathological processes (the human cancer colon tissue was obtained directly form the centre of tumor mass).^49^

Fig. 13 shows the data obtained for stomach tissues: normal and cancer by Raman spectroscopy and imaging including Raman maps and spectra characterising different areas of samples.

**Fig. 13.**
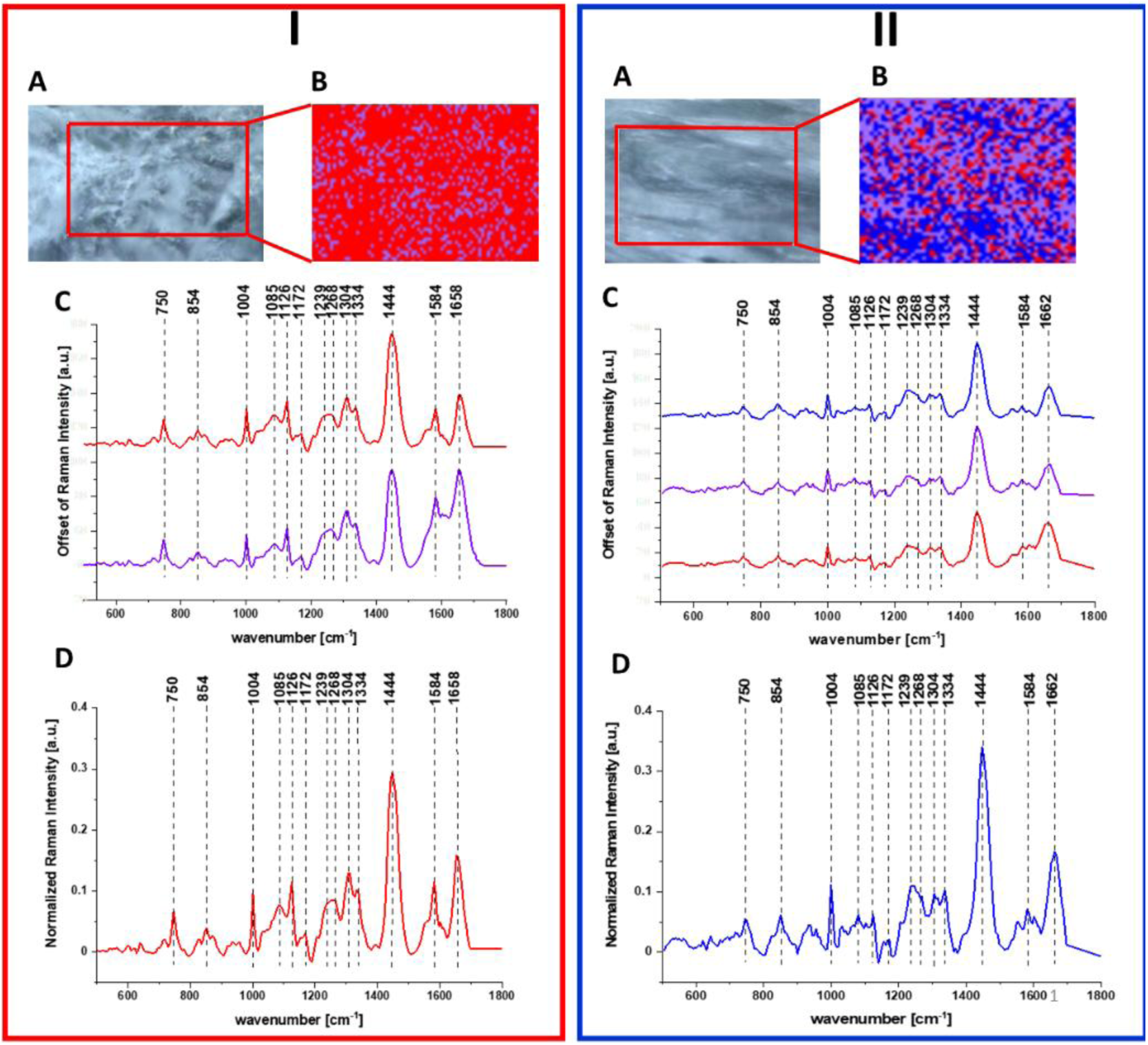
Raman image and spectral analysis of human gastric cancerous tissue (I) and human gastric normal tissue (II). The microscopy image (A), Raman image constructed by CA method (B), average Raman spectra typical of all clusters, colors of the spectra correspond to colors of clusters seen in B (C), and average (arithmetic mean) Raman spectrum for the entire area of analyzed tissue (D) for the human stomach cancer tissue, excitation wavelength 532 nm.

Once again based on Raman data presented in Fig. 13 we can characterize gastric human tissues by bands at 750, 854, 1004, 1085, 1126, 1172, 1239, 1268, 1304, 1334, 1444, 1584 and 1664 cm^−1^ which provide chemical information about the composition of cancerous and normal human gastric samples showed in a form of chemical images. Moreover, based on vibration features of tissues the difference in spectral profile between cancerous and normal tissue can be detected for nucleic acids, proteins and lipids, and the structural differences in regions within tissue areas can be presented in the form of clusters.

## Discussion

Gastrointestinal cancer is one of the most common cancer globally and oxidative stress is associated with the occurrence of carcinogenesis including CRC,^3^ and GC.^46^ Plant-derived compounds can find application as antioxidants, either by acting as ROS scavengers or by stimulating intracellular antioxidant enzymes.^50^

Vitamin C is a natural antioxidant commonly available in fruits and vegetables,^14^ or as a supplement.^12^ In our study, the effect of vitamin C on viability and composition of human colon cell CCD-18 Co exposed to oxidative stress was investigated and final conclusions confirm the anti-oxidant properties of this compound. CCD-18 Co cells are human fibroblast cells used in many studies as model normal colon cells, and different compounds such as linalool,^51^ Pt/MgO nanoparticles,^52^ or L-buthionine sulfoximine (BSO),^53^ were already reported to induce oxidative stress in CCD-18 Co cells. In our study, tBHP was used as an oxidative stress inducer, because this compound is also widely exploited to effectively induce oxidative stress *in vitro*.^54,55^

The effect of tBHP and vitamin C on viability of CCD-18 Co cells was determined by XTT tests. Viability of CCD-18 Co culture decreased in the presence of tBHP, but addition of vitamin C greatly improved viability of CCD-18 Co cells (scheme 1).

Stimulatory effect of vitamin C on metabolism of CCD-18 Co cells confirms results from a previous study.^48^ The cytotoxic effects of tBHP can be explain by loss of glutathione, lipid peroxidation and hemolysis. In addition, tBHP significantly increases the permeability of cell membranes and impairs the metabolic pathway leading to the synthesis of ATP and irreversible DNA damage. These effects were attributed to both radical and non-radical mechanisms.^56^

Over the course of the analysis of Raman spectra recorded for normal CCD-18 Co cells and CaCo-2 cancerous cells of the human colon, based on the intensity of the bands assigned to individual biochemical components, it can be noticed that vibrational spectroscopy effectively provides information on changes in the content of individual components depending on the type of analysed components.

Fig. 14 shows the differential spectrum of normal and cancerous cells (marked in green), the spectrum typical for the CaCo-2 cancer cell line (marked in red) and the spectrum typical for CCD-18 Co normal line (marked in blue) generated using Cluster Analysis for cells as a single cluster.

**Fig. 14.**
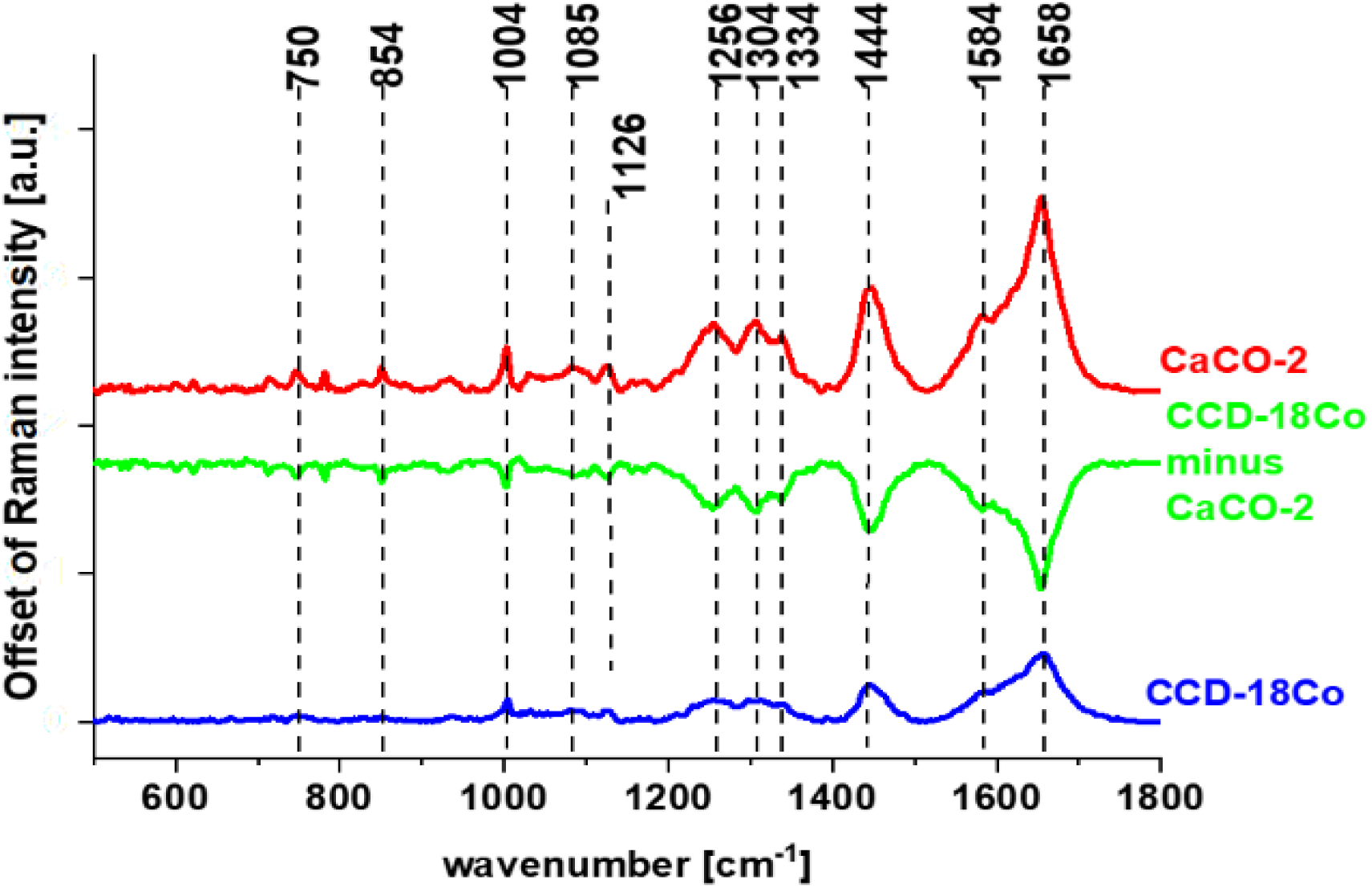
Average spectra for cancerous CaCo-2 (red) and normal CCD-18 Co (blue) human colon cells, and difference spectrum (CCD-18 Co minus CaCo-2), in the fingerprint region, excitation wavelength 532 nm.

The analysis of the differential spectrum presented in Fig. 14 shows that the most significant differences between normal and cancerous cells for human colon are observed for the following frequencies: 1004, 1256, 1304, 1334, 1444, 1584 and 1658 cm^−1^, which according to literature reports can be assigned to individual cell components such as: DNA, RNA, lipids, proteins or unsaturated fatty acids.^47^

Higher intensities of bands assigned to proline/hydroxyproline/tyrosine (854 cm^−1^), phenylalanine (1004 cm^−1^), Amide III (1256 cm^−1^), collagen (1334 cm^−1^) and Amide I (1658 cm^−1^) for CaCo-2 shows higher content of proteins for cancerous cells. It was concluded that increased protein content in CaCo-2 is, *inter alia*, due to increased nucleic acid (RNA/DNA) content in cancer cells,^44,45^ the band intensity for nucleic acid (750 cm^−1^, 1334 cm^−1^) is also higher in Raman spectral profile for CaCo-2. The band characterizing phosphorylated status of proteins (1584 cm^−1^) is also higher for cancer CaCo-2 cells than for normal CCD-18 Co cells, suggesting that the process of protein phosphorylation, involved in induction of cell proliferation, invasion, metastasis and inhibition of cell apoptosis,^57^ is up-regulated in cancer.

Fig. 15 shows average Raman spectra of CCD-18 Co cells, CCD-18 Co cells subjected to tBHP (50 µM) and CCD-18 Co cells subjected to tBHP upon supplementation with different vitamin C concentrations.

One can see from Fig. 15 that using Raman spectroscopy and imaging, the influence of the addition of a ROS generating agent and addition of vitamin C on the vibrational spectra of the tested cells can be assessed. Based on data presented in Fig. 15, which correspond to mean spectra determined for cells as a single cluster of pure cells, cells with the addition of tBHP, cells subjected to tBHP with the addition of vitamin C at various concentrations, the qualitative and quantitative analysis (discussed later) of cells can be performed. The comparison of spectral profiles shows differences for bands at 1004, 1078, 1258, 1444 and 1658 cm^−1^ assigned to proteins, DNA and lipids, confirming that tBHP and vitamin C treatment affect metabolism and structural composition of CCD-18 Co cells.

Analysis of Raman band intensities ratio of different bands within spectrum can provide further information regarding molecular characterization of cells. That’s why to find quantitative differences between analysed cells types we decided to focus on selected Raman bands typical for main chemical compounds: nucleic acid, proteins, lipids. Different ratios, for selected Raman band intensities corresponding to 1004/1078 (phenylalanine/nucleic acids and phospholipids), 1004/1258 (phenylalanine/amide III), 1004/1662 (phenylalanine/amide I proteins) and 1004/1444 (phenylalanine/lipids and proteins) were determined to monitor metabolic alterations in CCD-18 Co cells including treated with tBHP or tBHP and different concentrations of vitamin C.

The comparison of obtained results including data typical for cancer CaCo-2 cells is presented in Fig. 16. One has to noticed that, during the statistical data analysis the intensity of the peak at 1004 cm^−1^ was kept constant.

**Fig. 16.**
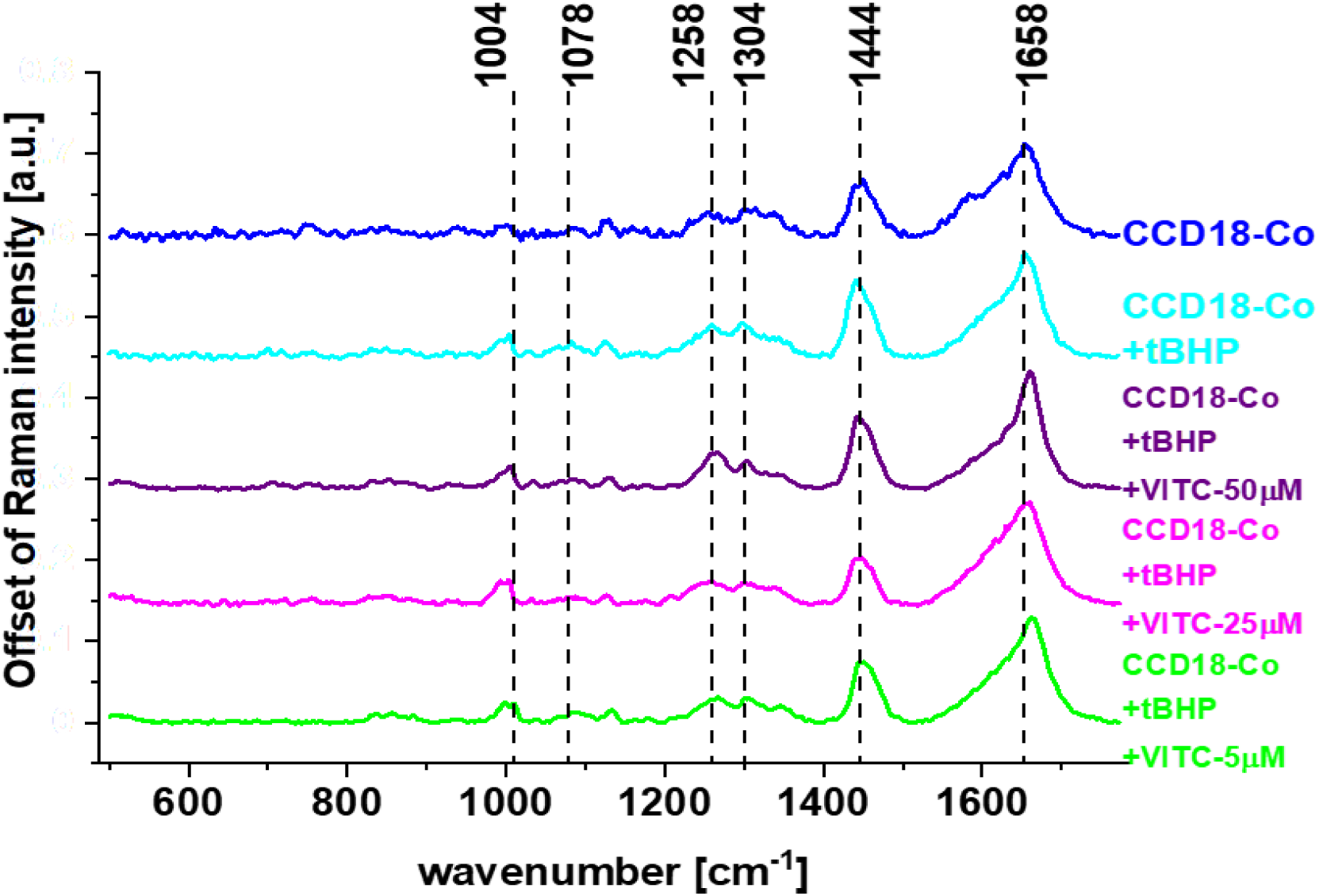
Average Raman spectra of CCD-18 Co cells, CCD-18 Co cells subjected to tBHP (50 µM) and CCD-18 Co cells subjected to tBHP upon supplementation with different vitamin C concentrations, excitation wavelength 532 nm.

One can see from Fig. 16 that values of investigated ratios changed with the change in concentrations of vitamin C used in experiments.

The first investigated ratio was I_1004_/_1078_, which is a ratio of phenylalanine to phosphodiester groups in nucleic acids and phospholipids.

Value of I_1004/1078_ was definitely higher for CCD-18 Co cells than for Caco-2 cells, which confirms that the amount of nucleic acids in cancer cells is higher than in normal cells (the intensity of the peak at 1004 cm^−1^ was constant and equal to 1.0). Treatment of CCD-18 Co cells with tBHP and with different vitamin C concentrations showed slight alterations in a value of I_1004_/_1078_, if compared to CCD-18 Co cells without any treatment. The higher value was observed upon supplementation with vitamin C with 5 µM concentration of ROS injured CCD-18 Co cells. It means that after generation of ROS the concentration of antioxidant is too low to protect cells from harmful effects of tBHP, and the protective effect can be seen for higher concentration of vitamin C. For the concentration of 50 µM the effect was the strongest. The lowest value for the ratio I_1004_/_1078_ was obtained for Caco-2 cancer cells. It was the expected results as the more intense peaks were observed also for cancer human colon tissue (see Fig. 12.I).

It has been shown that permanent changes in the genome of cells resulting from oxidative stress, is the first stage characteristic of the process of mutagenesis, carcinogenesis and cell aging. The appearance of a mutation in DNA is a critical step in the process of carcinogenesis. In various types of tumors an increase in the number of oxidative damage in DNA was observed.^58^ Our results presented and discussed above support this thesis. One has to remember also that in the stage of promoting carcinogenesis, ROS can induce proliferation or apoptosis of initiating cell clones. Under the influence of ROS, the concentration of Ca^2+^ ions increases significantly in cells, which can then activate some proto-oncogenes as well as protein kinase C and thus intensify cell proliferation and the stage of cancer promotion.^59^

The next ratio of interest I_1004_/_1258_ is a ratio of the intensity of bands related to the phenylalanine to amide III (N-H and C-H bend mode). A value of I_1004_/_1258_ increased for CCD-18 Co cells treated with tBHP confirming destructive effect generated by tBHP on proteins, but decreased sharply for higher vitamin C concentration (50 µM), which suggests the protective effect of this compound. The I_1004_/_1662_ ratio also correspond to proteins, and it’s value is related to phenylalanine to amide I (H-bonded C=O stretch mode). A value of this ratio did not differ with statistical significance between untreated CCD-18 Co cells and Caco-2 cells, but treatment of CCD-18 Co cells with tBHP increased I_1004_/_1662_ value, with the tendency for the graduate decrease while higher vitamin C concentrations were used.

At a high concentration of ROS, and at the same time with a reduced activity of proteolytic systems, oxidized proteins accumulate in the cell. Their presence has been detected in many tissues. It has been shown that oxidative stress and protein modifications caused by ROS play a role in both the aging process and the pathogenesis of many diseases including cancer.^60^ Our results obtained for ROS injured normal colon cells CCD-18 Co are consistent with this findings.

Another evaluated ratio, I_1004/1444_ comparing phenylalanine to lipids, showed that treatment of CCD-18 Co with tBHP increased a value of this ratio, while higher vitamin C concentrations affected the ratio differently. With further increase in I_1004_/_1444_ value for 25 µM vitamin C and a substantial decrease in I_1004_/_1444_ value for 50 µM vitamin C. The observed increasing tendency confirm once again the protective effect of vitamin C against ROS, based on the third type - lipids of main cells buildings compound: nucleic acids, proteins and lipids.

ROS can also destroyed lipids structures. In general, lipid peroxidation is the chain of reactions of oxidative degradation of lipids whit a final products in the form of lipid peroxides or lipid oxidation products e.g. reactive aldehydes, such as malondialdehyde (MDA) and 4-hydroxynonenal (HNE) (the second one being known as “second messenger of free radicals”).

Summarizing, analysis of Raman ratios (I_1004/1078_, I_1004/1258_ I_1004/1444_ and I_1004/1662_) show that tBHP and different vitamin C concentration treatment altered metabolism and chemical composition of human, normal colon CCD-18 Co cells. Spectroscopic data have proved also that RS can be effectively used for ROS injury and protective role of antioxidants tracking.

Using Raman spectroscopy, the influence of the addition of a vitamin E on the vibrational spectra of the tested gastric cells was also assessed.

Fig. 17 shows the average spectra for cancerous HTB-135 (red) and normal CRL-7869 (blue) human gastric cells, and difference spectrum (CRL-7869 minus HTB-135), in the fingerprint region. The figure shows mean spectra determined for cells as a single cluster of pure cells, and cells with the addition of vitamin E at various concentrations. The comparison of spectral profiles show differences for bands at 1004, 1085, 1256, 1444 and 1658 cm^−1^ assigned to proteins, DNA and lipids. The same obtained results are consistent with data presented for human colon samples.

**Fig. 17.**
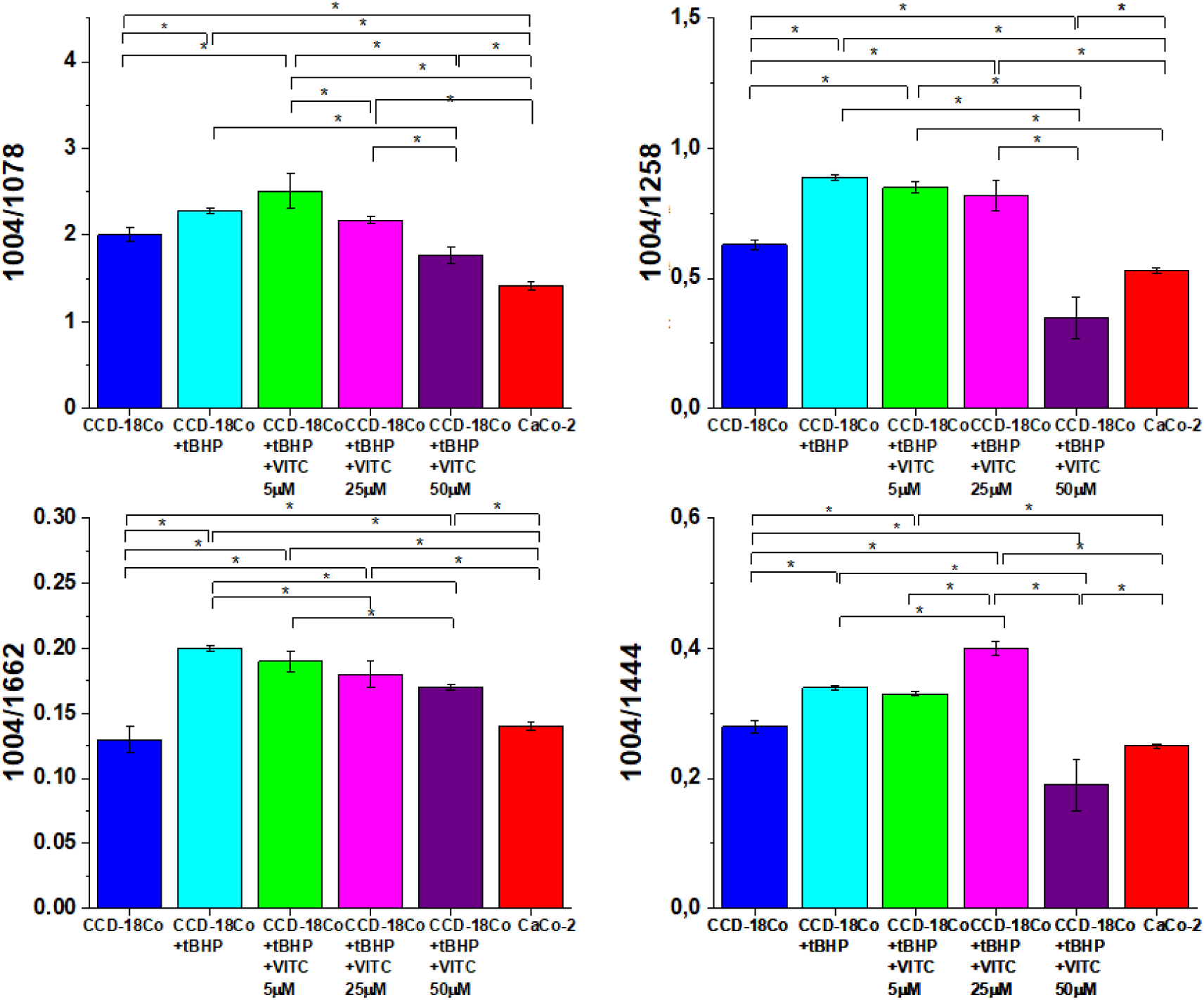
Raman band intensities ratios for selected Raman bands corresponding to 1004/1078, 1004/1258, 1004/1662, 1004/1444 in CCD-18 Co cells, in CCD-18 Co cells treated with tBHP and different concentrations of vitamin C. Plots show data obtained for 6 groups of normal human colon cells CCD18-Co: control group (labelled CCD-18-Co, blue), group supplemented with tBHP (labelled CCD-18-Co+tBHP, turquoise), group supplemented with tBHP and vitamin C in concentration of 5 µM (labelled CCD-18-Co+tBHP+VITC 5 µM, green), group. supplemented with tBHP and vitamin C in concentration of 25 µM (labelled CCD-18-Co+tBHP+VITC 25 µM, magenta), group supplemented with tBHP and vitamin C in concentration of 50 µM (labelled CCD-18-Co+tBHP+VITC 50 µM, violet), group of pure colon cancer cell measured in PBS (labelled CaCo-2, red), where the statistically significant results, based on ANOVA analysis have been marked with asterix. During the statistical data analysis the intensity of the peak at 1004 cm^−1^ was kept constant.

Under normal physiological conditions, the intracellular levels of ROS are steadily maintained to prevent cells from damage. Detoxification from ROS is facilitated by non-enzymatic molecules (i.e. glutathione, flavonoids and vitamins A, C and E) or through antioxidant enzymes which specifically scavenge different kinds of ROS.

In cancer cells high, abnormal levels of ROS can result from increased metabolic activity, mitochondrial dysfunction, peroxisome activity, increased cellular receptor signalling, oncogene activity, increased activity of oxidases, cyclooxygenases, lipoxygenases and thymidine phosphorylase, or through crosstalk with infiltrating immune cells. In mitochondria, ROS are produced as an inevitable by-product of oxidative phosphorylation. The electron transport chain encompasses complexes I-IV and ATP synthase on the mitochondrial inner membrane. Superoxide is generated at complexes I and III and released into the intermembrane space (approx. 80% of the generated superoxide) or the mitochondrial matrix (approx. 20%) and further the mitochondrial permeability transition pore in the outer membrane of the mitochondrion allows the leakage of superoxide into the cytoplasm. Growth factors and cytokines also stimulate the production of ROS to exert their diverse biological effects in cancer. Many cancers arise from sites of chronic irritation, infection, or inflammation. Recent data have expanded the concept that inflammation is a critical component of tumor progression.^61–63^ Macrophages induce the generation of ROS within tumor cells through secretion of various stimuli, such as TNFα.^64^ Oxidative stress-mediated signalling events have been reported to affect all characters of cancer cell behaviour.^65^ For instance, ROS in cancer are involved in cell cycle progression and proliferation, cell survival and apoptosis, energy metabolism, cell morphology, cell-cell adhesion, cell motility, angiogenesis and maintenance of tumor stemness.

Fig. 18 shows the average spectra for normal (CRL-7869) human gastric cells, cancer (HTB-135) human gastric cells, and human gastric cancer cells exposed to different vitamin E concentrations, in the fingerprint region. The frequencies differentiating untreated and supplemented by using vitamin E human cancer gastric cells are also highlighted.

**Fig. 18.**
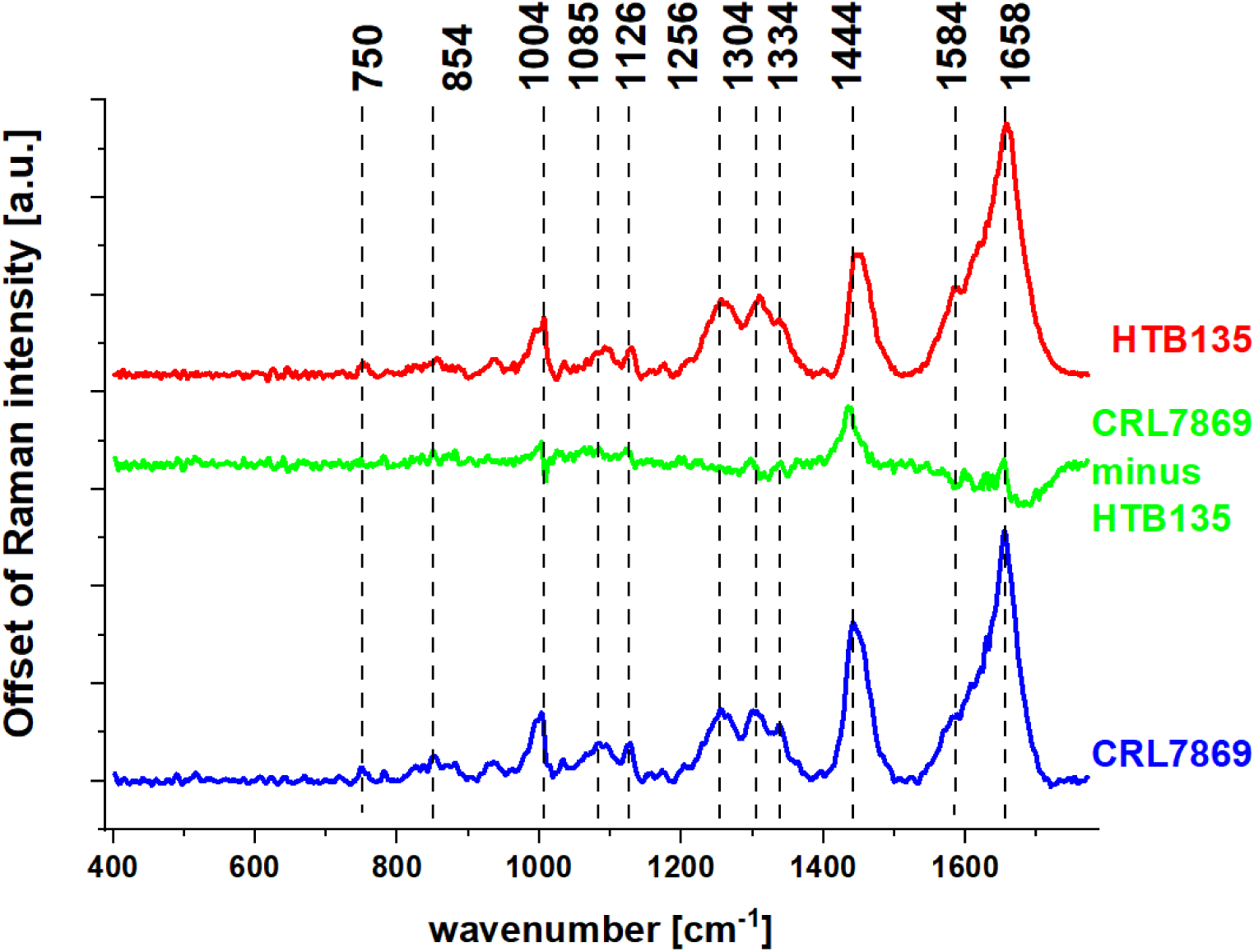
Average spectra for cancerous HTB-135 (red) and normal CRL-7869 (blue) human gastric cells, and difference spectrum (CRL-7869 minus HTB-135), in the fingerprint region, excitation wavelength 532 nm.

Ratios for selected Raman band intensities, used for characterisation of CCD-18 Co cells (I_1004/1078_, I_1004/1258_, I_1004/1658_ and I_1004/1444_), were also used to monitor metabolic alterations in HTB-135 cells, treated with different concentrations of vitamin E. These ratio values were also compared with other profiles: HTB-135 cells non-subjected to vitamin E exposure, and CRL-7869 cells (normal gastric cells). Values of investigated ratios as a function of vitamin E concentrations used in experiments are presented in Fig. 19.

**Fig. 19.**
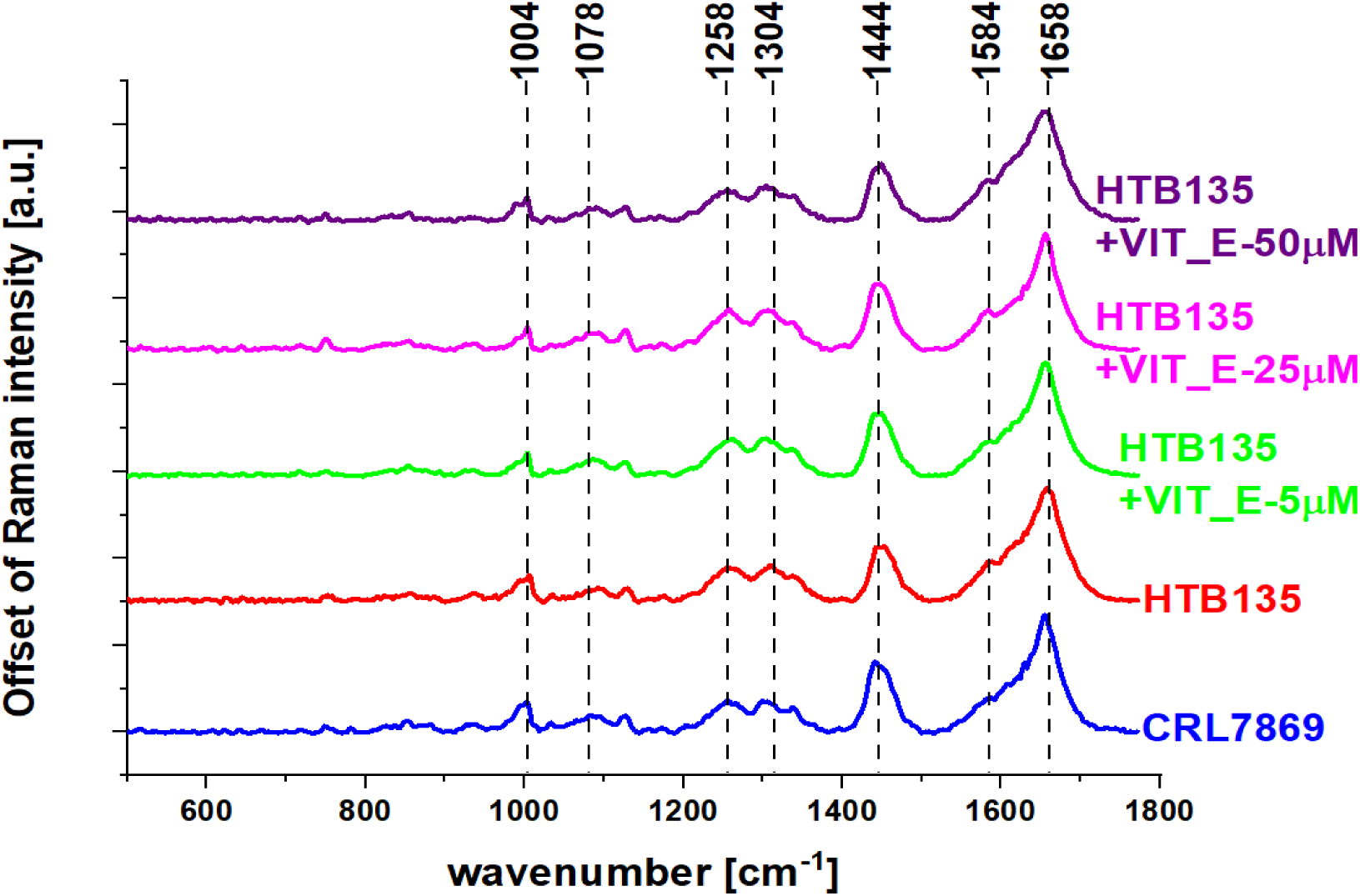
Average spectra for normal (CRL-7869) human gastric cells, cancer (HTB-135) human gastric cells, and cancer (HTB-135) human gastric cells exposed to different vitamin E concentrations, in the fingerprint region, excitation wavelength 532 nm.

**Fig. 20.**
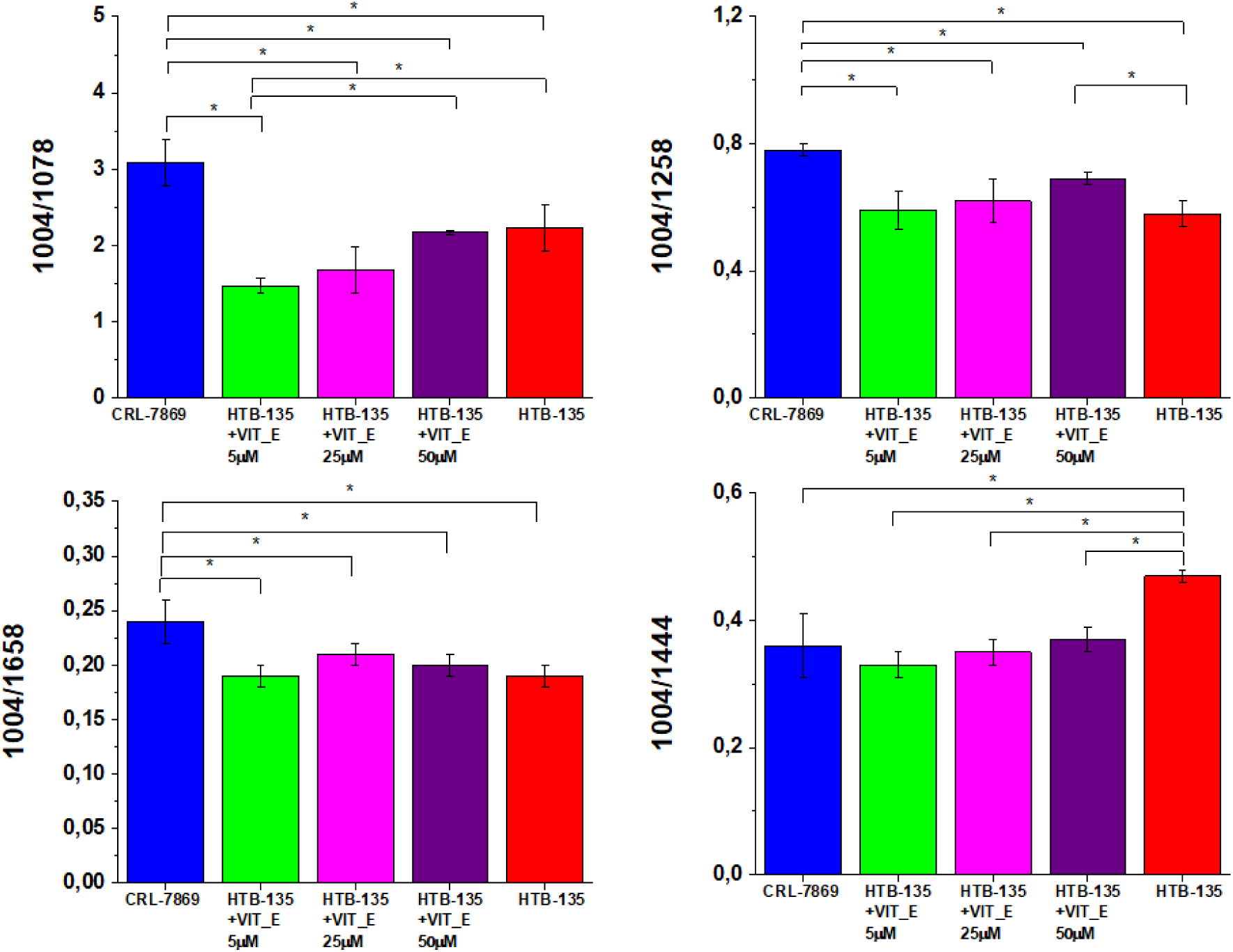
Raman band intensities ratios for selected Raman bands corresponding to 1004/1078, 1004/1258, 1004/1658 and 1004/1444, in HTB-135 cells, treated with different concentrations of vitamin E. Plots show data obtained for 5 groups of human gastric cancer cells HTB-135: control group (labelled HTB-135, red), group supplemented with vitamin E in concentration of 5 µM (labelled HTB-135 + VIT_E 5 µM, green), group supplemented with vitamin E in concentration of 25 µM (labelled HTB-135 + VIT_E 25 µM, magenta), group supplemented with vitamin E in concentration 50 µM (labelled HTB-135 + VIT_E 50 µM, violet), group of pure normal gastric cell (labelled CRL7869, blue) where the statistically significant results, based on ANOVA analysis have been marked with asterix. During the statistical data analysis the intensity of the peak at 1004 cm^−1^ was kept constant.

One can see from Fig. 19 that the value of I_1004/1078_ (phenylalanine/nucleic acids and phospholipids) was definitely higher for CRL-7869 cells than for HTB-135 cells profiles, this observation confirms the higher amount of nucleic acids in cancer gastric cells. Treatment of HTB-135 cells with vitamin E showed the decrease in a value of I_1004_/_1078_. When smaller vitamin E concentrations (5 µM) were tested, such a relation proved that low concentration of vitamin E cannot prevent the natural ROS damage typical for cancer cells and higher concentrations of antioxidant are needed to obtain the protective effect.

The ratio I_1004_/_1258_ (phenylalanine/amide III) is typical for tracking proteins. One can see from Fig. 19 that this ratio was higher for CRL-7869 cells than for HTB-135 cells profiles (the same the lower amount of proteins is typical for normal CRL-7869 gastric cells). The treatment of HTB-135 cells with 50 µM vitamin E forced I_1004_/_1258_ value up if compared to untreated HTB-135 cells, this confirm that vitamin E modulate the expressions of proteins in analysed cells. The higher the concentration of vitamin E the lower the signal 1258 cm^−1^ the higher the I_1004_/_1258_ ratio. Again, an I_1004_/_1658_ (phenylalanine/amide I proteins) ratio showed a higher value for CRL-7869 cells than for HTB-135 cells confirming the lower expressions of proteins in healthy cells.

In case of an I_1004/1444_ (phenylalanine/lipids), a value of this ratio was lower for CRL-7869 cells than for untreated HTB-135 cells, but treatment of HTB-135 cells with different vitamin E concentrations (5-50 µM) decreased I_1004/1444_ values to the levels of normal gastric cell profile which may suggest the protective role of vitamin E against natural oxidative processes typical for cancer samples.

Summarizing, the difference in values of I_1004/1078_, I_1004/1258_, I_1004/1658_ and I_1004/1444_ found between CRL-7869 and HTB-135 cells, showing profound metabolic changes distinguishing cancer and normal cells.

All presented results proved that metabolic regulations in cancer may involve genetic modifications and a change in metabolite pattern, with higher amino acid level, upregulated lipogenesis and upregulation of nucleotide metabolism and synthesis.^66^ Moreover, HTB-135 cells exposed to different vitamin E concentrations show further changes in I_1004/1078_, I_1004/1258_ and I_1004/1444_, indicating that vitamin E can affect intensity of Raman bands typical for nucleic acids, proteins and lipids in HTB-135 cells.

The connection between cancer risk and vitamin E has been presented in various scientific studies. The anticancer effects of vitamin E has been attributed mainly to its antioxidant, anti-inflammatory, anti-proliferative, anti-angiogenic, immune modulatory mechanisms and the inhibition of reductase enzyme - HMG CoA. Antioxidants such as vitamin E provide a protective effect through neutralization. This process occurs as a result of the donation of one of its own electrons. However, this action will not turn the antioxidants themselves into free radicals, as they are stable in both forms.^67^ Vitamin E reduces the activity of free radicals, preventing the escape of electrons, and thus directly participating in the formation of peroxide.^68^ The radical scavenging mechanism works by removing excess free radicals, and vitamin E is known to be one of the most abundant antioxidants, an important lipophilic antioxidant that scavenges radicals and effectively scavenges peroxide radicals.^69^ Moreover, vitamin E was reported to interrupt *de novo* sphingolipid pathway in a prostate cancer cell line,^33^ and modulate DNA synthesis in erythroleukemia, prostate and breast cancer cells.^35^

Moreover, increased mRNA and protein levels of PPARγ, involved in fatty acid uptake and transport, were reported in colon cancer cells (SW 480) exposed to vitamin E.^70^ Our study shows that vitamin E influences on nucleic acids, proteins and lipids in gastric cancer (HTB-135) cells, and all these changes can be tracking by Raman spectroscopy and imaging.

## Conclusions

In this work, Raman spectroscopy and imaging were used to evaluate the effect of vitamin C on molecular composition of human normal colon cells (CCD-18 Co) exposed to oxidative stress (tBHP), and the evaluate effect of vitamin E on biochemical composition of gastric cancer cells (HTB-135).

Raman spectroscopy and imaging successfully differentiated various types of colon and stomach cells based on their vibrational features. Moreover, Raman spectroscopy and imaging was successfully used to visualize organelles: nucleus, mitochondria, lipid structures, cytoplasm and cell membranes in single cells (CCD-18 Co, Caco-2, CRL-7869, HTB-135), according to vibrational bands in the fingerprint region of spectra.

Based on Raman band intensities attributed to proteins, nucleus acids and lipids as well as ratios 1004/1078, 1004/1258, 1004/1444, 1004/1658 calculated based on them, we have confirmed the protective role of vitamin C for cells in oxidative stress conditions for label-free and non-destructive spectroscopic method. For the same ratios we have confirmed the protective role of vitamin E for cancerous cells of human stomach.

Based on calculated ratios: 1004/1078, 1004/1258, 1004/1444, 1004/1658 we have proved also that the vitamin concentration effect on biochemical composition on CCD-18 Co and HTB-135 cell can be observed.

Our results proved that low doses of vitamins C and E can significantly reduce the risk of gastrointestinal cancer. More and greater clinical trials should be performed to define appropriate doses of vitamins in order to generate visible association between intake of antioxidants in the form of vitamin C and E and the risk of cancer of gastrointestinal tract.

Based on the obtained results, as well as the statistical analysis, we concluded that Raman spectroscopy enables the detection of cancerous changes in the human colon tissues based on the identification of characteristic vibrational bands of nucleic acids, proteins and lipids, including unsaturated fatty acids.

Moreover, gastric cancer and normal tissues were analysed by means of Raman spectroscopy and imaging, depicting the difference in the structure between cancerous and non-cancerous gastric tissue based on spectra profile with bands typical for proteins, nucleic acids and lipids in the fingerprint region.

## Author contributions

Conceptualization: BB-P; Funding acquisition: BB-P; Investigation: BB-P, KM, KB; Methodology: BB-P, Data analysis: BB-P, MK, KB, Writing – original draft: MK, BB-P, KB; Manuscript editing: KB, MK, BB-P. All authors reviewed and provide feedback on the manuscripts.

## References

1 C. G. Lengyel, B. Habeeb, S. Z. Khan, K. El Bairi, S. C. Altuna, S. Hussain, S. A. Mazher, D. Trapani and A. Petrillo, Gastrointest. Disord., 2020, 3, 1–22.

2 S. RL, M. KD, G. S. A, F. SA, B. LF, A. JC, C. A, S. RA and J. A, CA. Cancer J. Clin., 2020, 70, 145–164.

3 S. Lin, Y. Li, A. A. Zamyatnin, J. Werner and A. V. Bazhin, J. Cell. Physiol., 2018, 233, 5119–5132.

4 R. Li, M. Liang, X. Liang, L. Yang, M. Su and K. P. Lai, Front. Oncol., 2020, 10, 868.

5 F. A. Haggar and R. P. Boushey, Clin. Colon Rectal Surg., 2009, 22, 191–197.

6 J. Björk, EPMA J., 2010, 1, 513–521.

7 C. F, M. M, R. F, J. A, G. AG, A. K. S, B.-A. T, J. R, D. P, L. A and T. G, Anticancer Res., 2017, 37, 4759–4766.

8 I. IA and N. M, World J. Gastroenterol., 2017, 23, 5086–5096.

9 A. J. Bass, V. Thorsson, I. Shmulevich, S. M. Reynolds, M. Miller, B. Bernard, T. Hinoue, P. W. Laird, C. Curtis, H. Shen, D. J. Weisenberger, N. Schultz, R. Shen, N. Weinhold, D. P. Kelsen, R. Bowlby, A. Chu, K. Kasaian, A. J. Mungall, A. G. Robertson, P. Sipahimalani, A. D. Cherniack, G. Getz, Y. Liu, M. S. Noble, C. Pedamallu, C. Sougnez, A. Taylor-Weiner, R. Akbani, J. S. Lee, W. Liu, G. B. Mills, D. Yang, W. Zhang, A. Pantazi, M. Parfenov, M. Gulley, M. B. Piazuelo, B. G. Schneider, J. Kim, A. Boussioutas, M. Sheth, J. A. Demchok, C. S. Rabkin, J. E. Willis, S. Ng, K. Garman, D. G. Beer, A. Pennathur, B. J. Raphael, H. T. Wu, R. Odze, H. K. Kim, J. Bowen, K. M. Leraas, T. M. Lichtenberg, S. Weaver, M. McLellan, M. Wiznerowicz, R. Sakai, M. S. Lawrence, K. Cibulskis, L. Lichtenstein, S. Fisher, S. B. Gabriel, E. S. Lander, L. Ding, B. Niu, A. Ally, M. Balasundaram, I. Birol, D. Brooks, Y. S. N. Butterfield, R. Carlsen, J. Chu, E. Chuah, H. J. E. Chun, A. Clarke, N. Dhalla, R. Guin, R. A. Holt, S. J. M. Jones, D. Lee, H. A. Li, E. Lim, Y. Ma, M. A. Marra, M. Mayo, R. A. Moore, K. L. Mungall, K. M. Nip, J. E. Schein, A. Tam, N. Thiessen, R. Beroukhim, S. L. Carter, J. Cho, D. DiCara, S. Frazer, N. Gehlenborg, D. I. Heiman, J. Jung, J. Kim, P. Lin, M. Meyerson, A. I. Ojesina, C. S. Pedamallu, G. Saksena, S. E. Schumacher, P. Stojanov, B. Tabak, D. Voet, M. Rosenberg, T. I. Zack, H. Zhang, L. Zou, A. Protopopov, N. Santoso, S. Lee, J. Zhang, H. S. Mahadeshwar, J. Tang, X. Ren, S. Seth, L. Yang, A. W. Xu, X. Song, R. Xi, C. A. Bristow, A. Hadjipanayis, J. Seidman, L. Chin, P. J. Park, R. Kucherlapati, S. Ling, A. Rao, J. N. Weinstein, S. B. Kim, Y. Lu, M. S. Bootwalla, P. H. Lai, T. Triche, D. J. Van Den Berg, S. B. Baylin, J. G. Herman, B. A. Murray, B. A. Askoy, G. Ciriello, G. Dresdner, J. Gao, B. Gross, A. Jacobsen, W. Lee, R. Ramirez, C. Sander, Y. Senbabaoglu, R. Sinha, S. O. Sumer, Y. Sun, L. Iype, R. W. Kramer, R. Kreisberg, H. Rovira, N. Tasman, D. Haussler, J. M. Stuart, R. G. W. Verhaak, M. D. M. Leiserson, B. S. Taylor, A. D. Black, J. A. Carney, J. M. Gastier-Foster, C. Helsel, C. McAllister, N. C. Ramirez, T. R. Tabler, L. Wise, E. Zmuda, R. Penny, D. Crain, J. Gardner, K. Lau, E. Curely, D. Mallery, S. Morris, J. Paulauskis, T. Shelton, C. Shelton, M. Sherman, C. Benz, J. H. Lee, K. Fedosenko, G. Manikhas, O. Potapova, O. Voronina, D. Belyaev, O. Dolzhansky, W. K. Rathmell, J. Brzezinski, M. Ibbs, K. Korski, W. Kycler, R. Lazniak, E. Leporowska, A. Mackiewicz, D. Murawa, P. Murawa, A. Spychala, W. M. Suchorska, H. Tatka, M. Teresiak, R. Abdel-Misih, J. Bennett, J. Brown, M. Iacocca, B. Rabeno, S. Y. Kwon, Kemkes, E. Curley, I. Alexopoulou, J. Engel, J. Bartlett, M. Albert, D. Y. Park, R. Dhir, J. Luketich, R. Landreneau, Y. Y. Janjigian, E. Cho, M. Ladanyi, L. Tang, S. J. McCall, Y. S. Park, J. H. Cheong, J. Ajani, M. C. Camargo, S. Alonso, B. Ayala, M. A. Jensen, T. Pihl, R. Raman, J. Walton, Y. Wan, G. Eley, K. R. M. Shaw, R. Tarnuzzer, Z. Wang, L. Yang, J. C. Zenklusen, T. Davidsen, C. M. Hutter, H. J. Sofia, R. Burton, S. Chudamani and J. Liu, Nat. 2014 5137517, 2014, 513, 202–209.

10 R. P and B. A, Prz. Gastroenterol., 2019, 14, 26–38.

11 R. Sitarz, M. Skierucha, J. Mielko, G. J. A. Offerhaus, R. Maciejewski and W. P. Polkowski, Cancer Manag. Res., 2018, 10, 248.

12 N. Yamamichi, C. Hirano, Y. Takahashi, C. Minatsuki, C. Nakayama, R. Matsuda, T. Shimamoto, C. Takeuchi, S. Kodashima, S. Ono, Y. Tsuji, M. Fujishiro, R. Wada, T. Mitsushima and K. Koike, Gastric Cancer, 2016, 19, 670–675.

13 K. GH, L. PS, B. SJ and H. JH, Gastrointest. Endosc., 2016, 84, 18–28.

14 C. Hamashima, World J. Gastroenterol., 2016, 22, 6392.

15 C. N, M. M, T.-Z. E, B. AN and B. M, Gut, 2020, 69, 2244–2255.

16 G. M. Cragg and J. M. Pezzuto, Med. Princ. Pract., 2016, 25, 59.

17 D. AL and V. P, Microb. Biotechnol., 2011, 4, 687–699.

18 J. Wang and Y.-F. Jiang, World J. Exp. Med., 2012, 2, 57.

19 J. A, T. A, V. A and J. SK, Curr. Mol. Med., 2017, 17, 321–340.

20 Y. Zhang, W. Zhou, J. Yan, M. Liu, Y. Zhou, X. Shen, Y. Ma, X. Feng, J. Yang and G. Li, Mol. 2018, Vol. 23, Page 1484, 2018, 23, 1484.

21 S. Chambial, S. Dwivedi, K. K. Shukla, P. J. John and P. Sharma, Indian J. Clin. Biochem., 2013, 28, 328.

22 P. G and H. HP, Adv. Biochem. Eng. Biotechnol., 2014, 143, 143–188.

23 X.-Y. Chen, Y. Chen, C.-J. Qu, Z.-H. Pan, Y. Qin, X. Zhang, W.-J. Liu, D.-F. Li and Q. Zheng, Oncol. Lett., 2019, 18, 3886.

24 A. Nagappan, H. S. Park, K. Il Park, J. A. Kim, G. E. Hong, S. R. Kang, J. Zhang, E. H. Kim, W. S. Lee, C. K. Won and G. S. Kim, BMC Biochem., 2013, 14, 24.

25 W. L, L. X, L. C, H. Y, X. P, L.-D. LH, D. T, P. C, G. R, Q. H, T. C and B. P, Oxid. Med. Cell. Longev.,, DOI:10.1155/2017/3481710.

26 J. Du, S. M. Martin, M. Levine, B. A. Wagner, G. R. Buettner, S. H. Wang, A. F. Taghiyev, C. Du, C. M. Knudson and J. J. Cullen, Clin. Cancer Res., 2010, 16, 509–520.

27 J. Eun Kim, K. Jae Seung and L. Wang Jae, IMMUNE Netw., 2012, 12, 189–195.

28 T. Y, S. M, S. K, H. H, S. Y and K. S, Biochem. Biophys. Res. Commun., 2010, 394, 249–253.

29 G. G. Martinovich, E. N. Golubeva, I. V. Martinovich and S. N. Cherenkevich, J. Biophys., 2012, 2012, 1–6.

30 P. Mathur, Z. Ding, T. Saldeen and J. L. Mehta, Clin. Cardiol., 2015, 38, 576.

31 S. Péter, A. Friedel, F. F. Roos, A. Wyss, M. Eggersdorfer, K. Hoffmann and P. Weber, https://doi.org/10.1024/0300-9831/a000281, 2016, 85, 261–281.

32 F. Shahidi and A. C. De Camargo, Int. J. Mol. Sci., 2016, 17, 1745.

33 J. Q, W. J, F. H, S. JD and A. BN, Proc. Natl. Acad. Sci. U. S. A., 2004, 101, 17825–17830.

34 Z. R, A. R. F and Z. RB, Integr. Cancer Ther., 2017, 16, 414–425.

35 G. Sigounas, A. Anagnostou and M. Steiner, Nutr. Cancer, 1997, 28, 30–35.

36 S. Lim, J. Y. Lee, W. H. Jung, E. H. Lim, M. K. Joo, B. J. Lee, J.-J. Park, J. S. Kim, Y.-T. Bak, S. W. Jung and S. W. Lee, Korean J. Helicobacter Up. Gastrointest. Res., 2011, 11, 175.

37 N. J, S. I, W. T, W. C and B. UT, FEBS Lett., 1999, 445, 295–300.

38 D. S, B. KN, B. SL, P. O, K. S, N. VM, S. F, N. R and C. IJ, Med. Phys.,, DOI:10.1118/1.4870981.

39 A. GW, K. SK, H. C, B. B, T. M, A. Z, E. A, M. KC and B. MA, Cancer Metastasis Rev., 2018, 37, 691–717.

40 M. G. Ramírez-Elías and F. J. González, Raman Spectrosc.,, DOI:10.5772/INTECHOPEN.72933.

41 C. S, Z. S and Y. S, J. Healthc. Eng.,, DOI:10.1155/2018/8619342.

42 A. Imiela, J. Surmacki and H. Abramczyk, J. Mol. Struct., 2020, 1217, 128381.

43 H. Abramczyk, J. Surmacki, M. Kopeć, A. K. Olejnik, A. Kaufman-Szymczyk and K. Fabianowska-Majewska, Analyst, 2016, 141, 5646–5658.

44 B. Brozek-Pluska, J. Mol. Struct.,, DOI:10.1016/j.molstruc.2020.128524.

45 B. Brozek-Pluska, A. Jarota, R. Kania and H. Abramczyk, Molecules, 2020, 25, 2688.

46 B. LD, den H. G, E. PB and C. SE, Cell. Mol. Gastroenterol. Hepatol., 2017, 3, 316–322.

47 Z. Movasaghi, S. Rehman and I. U. Rehman, Appl. Spectrosc. Rev., 2007, 42, 493–541.

48 K. Beton and B. Brozek-Pluska, Int. J. Mol. Sci. 2021, Vol. 22, Page 6928, 2021, 22, 6928.

49 B. Brozek-Pluska, J. Musial, R. Kordek and H. Abramczyk, Int. J. Mol. Sci. 2019, Vol. 20, Page 3398, 2019, 20, 3398.

50 M. J. Vallejo, L. Salazar and M. Grijalva, Oxid. Med. Cell. Longev.,, DOI:10.1155/2017/4586068.

51 K. Iwasaki, Y.-W. Zheng, S. Murata, H. Ito, K. Nakayama, T. Kurokawa, N. Sano, T. Nowatari, M. O. Villareal, Y. N. Nagano, H. Isoda, H. Matsui and N. Ohkohchi, http://www.wjgnet.com/, 2016, 22, 9765–9774.

52 M. Q. Al-Fahdawi, F. A. J. Al-Doghachi, Q. K. Abdullah, R. T. Hammad, A. Rasedee, W. N. Ibrahim, H. A. Alshwyeh, A. A. Alosaimi, S. K. Aldosary, E. E. M. Eid, R. Rosli, Y. H. Taufiq-Yap, N. A. Al-Haj and M. S. Al-Qubaisi, Biomed. Pharmacother., 2021, 138, 111483.

53 T. C. Ooi, K. M. Chan and R. Sharif, Biol. Trace Elem. Res. 2020 1982, 2020, 198, 464–471.

54 C. Wenz, D. Faust, B. Linz, C. Turmann, T. Nikolova and C. Dietrich, Arch. Toxicol. 2019 935, 2019, 93, 1265–1279.

55 C. M, G. H, Y. Y, L. B, S. L, D. W, Z. J and L. J, Food Chem. Toxicol., 2010, 48, 2980–2987.

56 S. Hix, M. B. Kadiiska, R. P. Mason and O. Augusto, Chem. Res. Toxicol., 2000, 13, 1056–1064.

57 X. Liu, Y. Zhang, Y. Wang, M. Yang, F. Hong and S. Yang, Biomol. 2021, Vol. 11, Page 1009, 2021, 11, 1009.

58 D. DA and P. M, Free Radic. Biol. Med., 2008, 44, 1873–1886.

59 D. D and J. AF, Eur. J. Cancer, 1996, 32A, 30–38.

60 M. B. Ponczek and B. Wachowicz, Postepy Biochem., 2005, 51, 140–145.

61 H. Xiao and C. S. Yang, Int. J. Cancer, 2008, 123, 983–990.

62 B. BM, Curr. Opin. Hematol., 1995, 2, 55–60.

63 S. AW and S. KP, Ann. N. Y. Acad. Sci., 1997, 832, 215–222.

64 K. Rabold, A. Aschenbrenner, C. Thiele, C. K. Boahen, A. Schiltmans, J. W. A. Smit, J. L. Schultze, M. G. Netea, G. J. Adema and R. T. Netea-Maier, J. Immunother. Cancer,, DOI:10.1136/JITC-2020-000638.

65 A. Gupta, S. F. Rosenberger and G. T. Bowden, Carcinogenesis, 1999, 20, 2063–2073.

66 X. S and Z. L, Int. J. Oncol., 2017, 51, 5–17.

67 L. A. Pham-Huy, H. He and C. Pham-Huy, Int. J. Biomed. Sci., 2008, 4, 96.

68 C. K. Chow, Biol. Sicals Recept., 2001, 10, 112–124.

69 N. E, Free Radic. Biol. Med., 2014, 66, 3–12.

70 S. E. Campbell, W. L. Stone, S. G. Whaley, M. Qui and K. Krishnan, BMC Cancer 2003 31, 2003, 3, 1–13.

